# Circular-Linear Copulae for Animal Movement Data

**DOI:** 10.1101/2021.07.14.452404

**Authors:** Florian H. Hodel, John R. Fieberg

## Abstract

1. Animal movement is often modeled in discrete time, formulated in terms of steps taken between successive locations at regular time intervals. Steps are characterized by the distance between successive locations (*step-lengths*) and changes in direction (*turn angles*). Animals commonly exhibit a mix of directed movements with large step lengths and turn angles near 0 when traveling between habitat patches and more wandering movements with small step lengths and uniform turn angles when foraging. Thus, step-lengths and turn angles will typically be cross-correlated.
2. Most models of animal movement assume that step-lengths and turn angles are independent, likely due to a lack of available alternatives. Here, we show how the method of copulae can be used to fit multivariate distributions that allow for correlated step lengths and turn angles.
3. We describe several newly developed copulae appropriate for modeling animal movement data and fit these distributions to data collected on fishers (*Pekania pennanti*). The copulae are able to capture the inherent correlation in the data and provide a better fit than a model that assumes independence. Further, we demonstrate via simulation that this correlation can impact movement patterns (e.g. rates of dispersion overtime).
4. We see many opportunities to extend this framework (e.g. to consider autocorrelation in step attributes) and to integrate it into existing frameworks for modeling animal movement and habitat selection. For example, copula could be used to more accurately sample available locations when conducting habitat-selection analyses.

## 1 INTRODUCTION

Advances in sensor technologies have led to the proliferation of animal location data at fine temporal and spatial scales (Kays et al., 2015), which in turn has led to the development of a plethora of new methods and software for modeling animal movements (see e.g., Hooten et al. 2017 for an overview of statistical methods and Joo et al. 2020 for a review of R packages for modeling animal movement). Although movement is inherently a continuous process, we usually observe it at discrete points in time, and the time interval between successive locations is also often, though not always, constant (e.g. Global Positioning Systems are typically programmed to collect data at fixed intervals). Thus, it is often easier to conceptualize movement in discrete time, with movements deconstructed into a series of “steps” between successive locations (McClintock et al., 2014). These steps can be characterized by the distance between locations (*step length*) and the change in direction from the previous step (*turn angle*). When modeling movement in discrete-time, it is important to consider the following characteristics of observed movement trajectories: 1) over short time scales, animals will tend to move in a consistent direction; 2) left- and right-turns are equally likely in the absence of environmental heterogeneity; and 3) animals will tend to move large distances with little change in direction when traveling between habitat patches and short distances with many changes in direction when foraging. These characteristics, in turn, suggest that: 1) the distribution of turn angles is likely to have a mode at 0 (indicative of directional persistence); 2) the distribution of turn angles should be symmetric around 0; and 3) step-lengths and turn angles are likely to be correlated. Commonly used circular distributions (e.g., wrapped Cauchy, von Mises) allow for symmetric angles about a mode of 0 and mixtures of circular distributions can accommodate multimodal angular densities. Thus, common circular distributions satisfy the first two conditions noted above. The third characteristic is more challenging to address. Correlation between direction and speed (equivalent to distance when observations are equally spaced in time) is an inherent property of continuous-time models (Johnson et al., 2008; McClintock et al., 2014; Calabrese et al., 2016; Gurarie et al., 2017) and discrete-time models formulated in terms of a velocity process (i.e., changes in position between successive time points; Jonsen et al., 2005). Yet, it is more common to parameterize discrete-time models using statistically independent step lengths and turn angles (Morales et al., 2004; McClintock et al., 2012, 2014). The main reason for assuming independence has been a lack of available analytical solutions that allow for proper modeling of this correlation. The broad goal of this paper is to address this limitation.

Correlation is easily accommodated when data are distributed according to a multivariate normal distribution, but our interest lies in modeling correlation between a linear (i.e. non-periodic) variable that can only take on positive values (step length), modeled using e.g. an exponential or gamma distribution, and a circular variable (turn angle), which is often modeled using a von Mises or other circular distribution (Morales et al., 2004; Avgar et al., 2016; Signer et al., 2019). Here, we show how copulae (Nelsen, 2006), cumulative distribution functions (CDFs) with uniform marginals on the interval [0,1], can be used to develop multivariate distributions that allow for correlation between step lengths and turn angles while preserving these intended marginal distributions. Specifically, we introduce several new circular-linear copulae with properties appropriate for modeling movement in discrete time together with corresponding methods for estimating parameters and simulating data. Functions for implementing these methods are available in our open-source `R`-package `cylcop` (R Core Team, 2019; Hodel and Fieberg, 2021), which can be seen as an extension of the `copula` package (Hofert et al., 2020; Jun Yan, 2007; Ivan Kojadinovic and Jun Yan, 2010; Marius Hofert and Martin Mächler, 2011; Hofert et al., 2018) and is available on GitHub at `floo66/cylcop`. Finally, we have also developed an app which allows the reader to interactively visualize all copulae introduced in this paper. The app can be accessed at https://cylcop.shinyapps.io/cylcop-graphs/.

We begin by providing a motivating example using GPS data collected on fishers (*Pekania pennanti*) from (LaPoint et al., 2013a,b). We expect most ecologists will be unfamiliar with copulae, so we follow this example with a short introduction to the basic theory (section 3) before presenting new methods for producing circular-linear copulae suitable for animal movement data (section 4). In section 5, we describe methods for estimating parameters of copulae. In section 6, we apply the introduced copulae and estimation methods to the fisher data. Finally, in the last section, we discuss possible future research directions.

## 2 MOTIVATING EXAMPLE

To illustrate typical correlation in movement data, we will consider tracking data from fishers (*Pekania pennanti*) available through Movebank (Brown et al., 2012; LaPoint et al., 2013a,b). The data, displayed in Figure 1a, consist of a total of 4350 step lengths and turn angles of 6 individuals, resampled to an interval of 10 minutes with a tolerance of 1 minute using the `track_resample` function in the `R`-package `amt` (Signer et al., 2019). The marginal density of the turn angles is symmetric around 0 and has modes at 0 and 180 degrees. The mode at 180 degrees could be due to the animals’ movements being influenced by linear features in the their environment (Mckenzie et al., 2012), or GPS measurement errors (Hurford, 2009). To evaluate measurement error as a potential explanation, we visualized the distribution of turn angles after excluding steps shorter than 25 m (Hurford, 2009), but this did not change the general shape of the distribution. Therefore, rather than exclude short steps, we will consider models that can accommodate multiple modes. The correlation between turn angles and step lengths can be seen in panel b of Figure 1. For the lower quintiles of step lengths, the circular medians and modes of the corresponding turn angles are close to *π*, whereas in the fourth and fifth quintiles, the medians and modes of the corresponding turn angles are close to 0. Thus, in this data set, step lengths and turn angles are clearly not independent.

**FIGURE 1.**
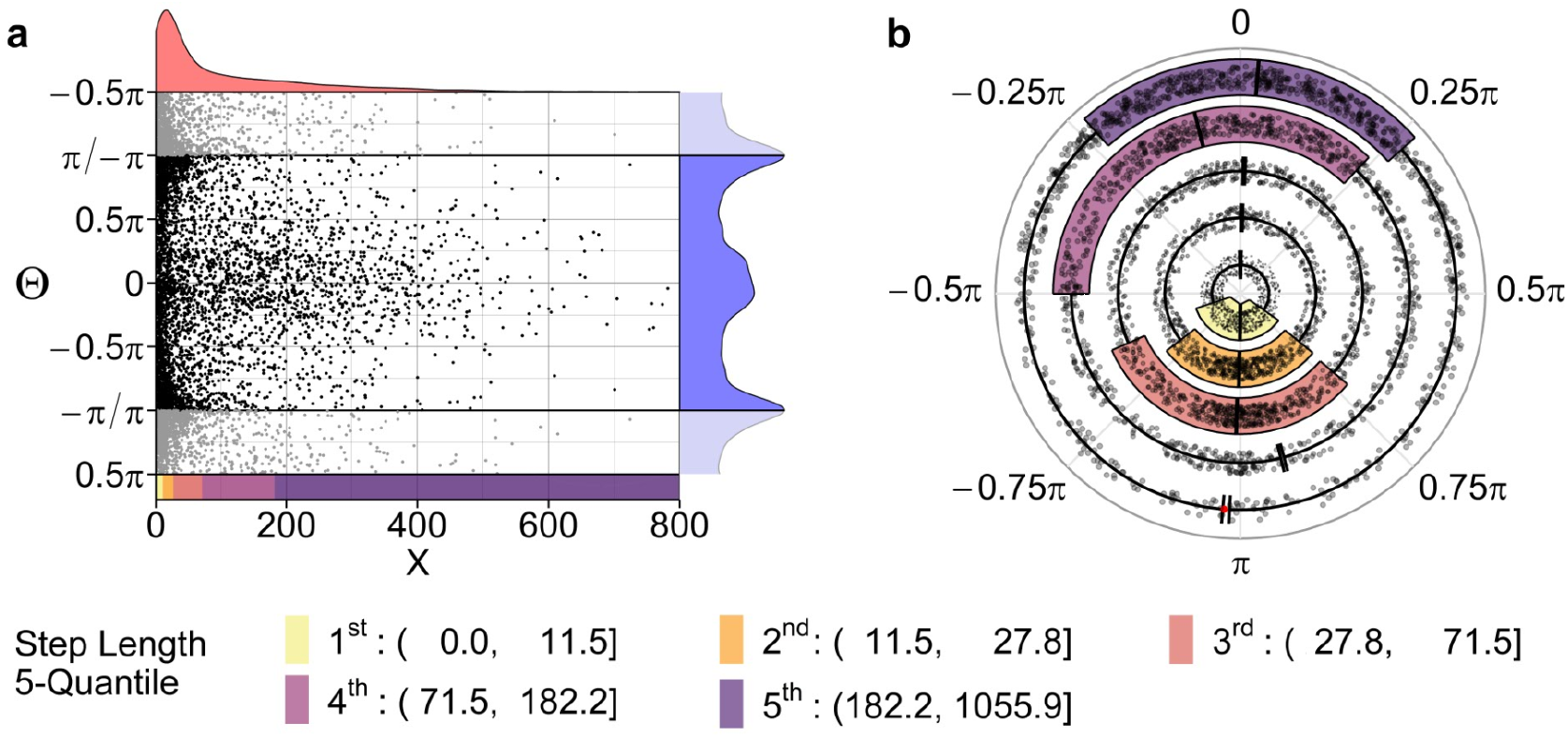
Left: Scatter plot of step lengths and turn angles of 6 fishers from (LaPoint et al., 2013a, b). Kernel density approximations of the marginal densities are plotted in red and blue next to the axes. The step lengths are separated into 5 quantiles as marked by the colorbar below the X-axis. Right: Circular box plots of the same data. For each of the 5 step length quantiles, a box plot of the corresponding angles is shown. The sample medians are defined as the center of the shortest arc that connects all points, in other words, the angle around which the spread of the data is minimal. Outliers, i.e. values outside 1.5 times the inter quartile range are marked in red.

## 3 COPULA

A bivariate copula *C*(*u, v*) is the CDF of a pair of random variables (*U, V*) with uniform marginals and its domain restricted to the unit square, (*u, v*) ∈ [0,1]^2^. Although we focus on bivariate copulae (referred to hereafter as “copulae”), extensions to more than 2 dimensions are straightforward. The main properties of copulae can be directly derived from well-known properties of general CDFs:

1. Any CDF must vanish at the lower limits and reach 1 at the upper limits of the domains of its random variables, *C*(*u*, 0) = *C*(0, *v*) = 0 and *C*(1, 1) = 1, ∀*u, v* ∈ [0, 1].
2. The CDF of a uniform random variable (and hence the marginal CDFs of a copula) is equal to the values of the random variable, *C*(*u*, 1) = *u* and *C*(1, *v*) = *v*, ∀*u, v* ∈ [0,1].
3. A bivariate CDF of a pair of random variables (*U, V*) cannot decrease with either increasing *u* or *v*, i.e. the probability of (*U, V*) taking a value from any subset of the entire domain must be non-negative.

To illustrate the usefulness of copulae, let *X* and *Y* be any two random variables with CDFs *F_X_*(*x*) = P(*X* ≤ *x*) and *F_Y_*(*y*) = P(*Y* ≤ *y*). Sklar’s theorem (Sklar, 1959) states that with marginal CDFs as arguments, a copula returns the CDF of a multivariate distribution, *F_X,Y_*(*x, y*).

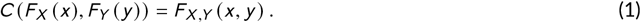

Thus, we can describe the joint distribution of *X* and *Y* using their specified marginal distributions together with a copula. All information about correlation between *X* and *Y* is “encoded” in the copula, and different copulae give different joint distributions for two specified marginal distributions. Consequently, when modeling data, we can focus separately on describing: a) the marginal distributions of *X* and *Y*; and b) modeling their dependence structure.

To further demonstrate, note that equation 1 can be used to define new copulae from a given joint distribution and its marginals, since, using the probability integral transform, we can rewrite it as 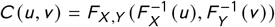, where *U* and *V* are again uniform random variables. As an example, consider the joint CDF Φ_*ρ*_ of a bivariate normal distribution with correlation *ρ* and both variances equal to one. Its marginal distributions are both standard normal distributions with CDF Φ. From this, we can generate a Gaussian or normal copula

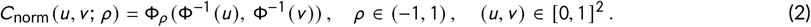

Now, let *X* and *Y* be some (non-standard) normally distributed random variables with CDFs *F_X_*(*x*) and *F_Y_*(*y*). One possible joint distribution with marginals *F_X_*(*x*) and *F_Y_*(*y*) is then

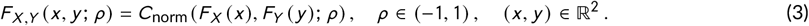

It can be easily shown that *F_X,Y_*(*x, y; ρ*) is a bivariate normal distribution with correlation *ρ* (see Fig. 2c). However, if we had instead chosen *X* to follow a gamma and *Y* an exponential distribution, we could still obtain a valid joint CDF *F_X,Y_*(*x, y; ρ*) from a Gaussian copula, but the resulting joint distribution would then not be bivariate normal (see Fig. 2e). Although the parameter *ρ* would continue to capture the correlation between *X* and *Y*, the exact Pearson correlation coefficient between *X* and *Y* would not be *ρ*, but a function of *ρ*. There are many other ways of generating copulae besides using known joint distributions, and as long as the obtained function fulfills the three conditions described above, it is a copula.

**FIGURE 2.**
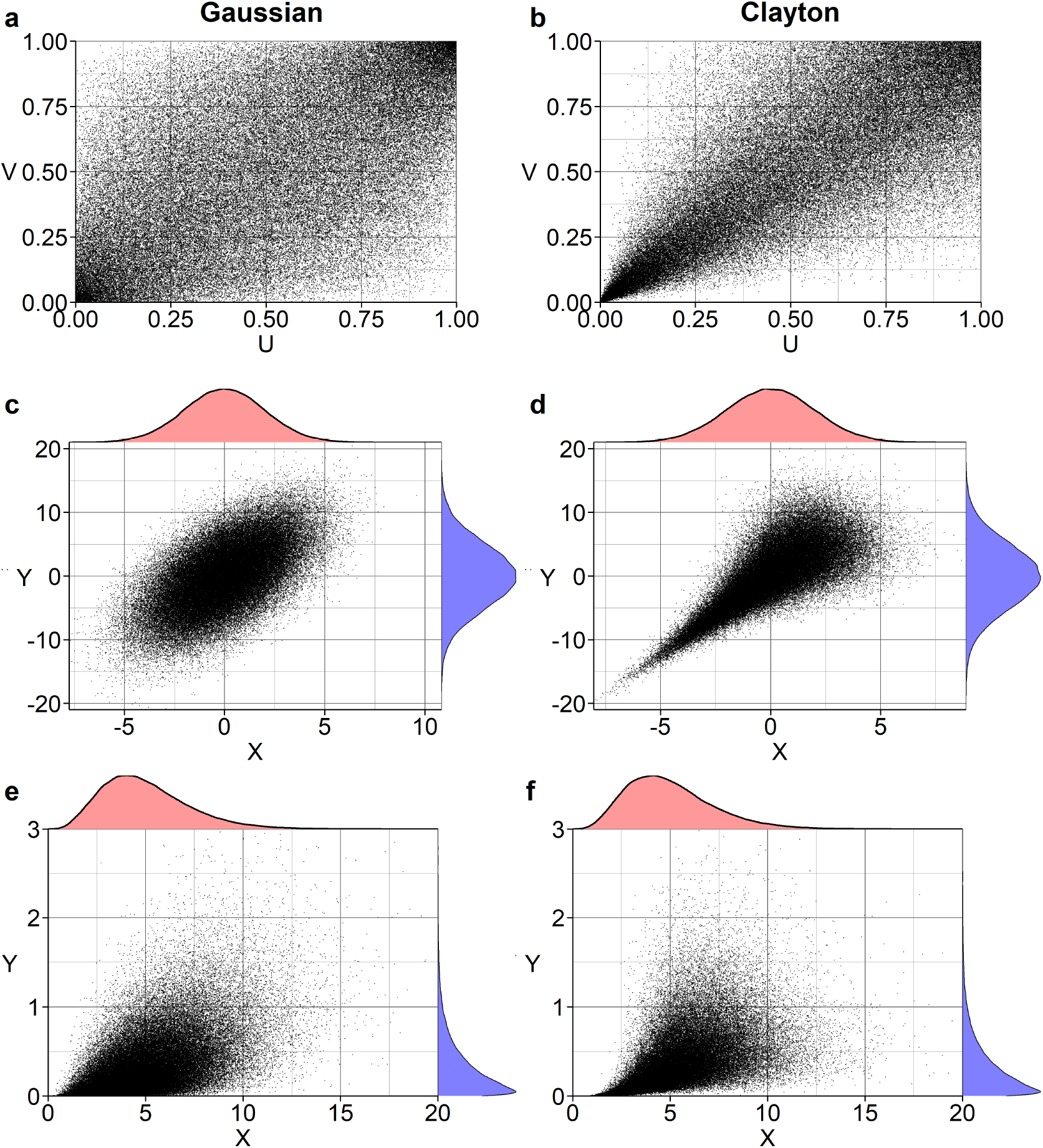
Samples from copulae and corresponding joint distributions with non-uniform margins. Left column: Gaussian copula with *ρ* = 0.6. Right column: Clayton copula with *α* = 3. First row: samples drawn from the copulae. Second row: samples drawn from the joint distribution obtained with the corresponding copula and normal margins with means 0 and standard deviations 2 and 5. Third row: samples drawn from the joint distribution obtained with the corresponding copula, a marginal gamma distribution with shape=5 and scale=1 (X-direction) and marginal exponential distribution with rate=3 (Y-direction).

The probability density function (PDF), *f_X,Y_*(*x, y*), of a joint distribution can also be obtained with the copula

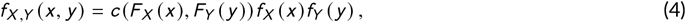

where *f_X_*(*x*) and *f_Y_*(*y*) denote the marginal PDFs of *X* and *Y*, respectively, and *c*(*u, v*) is the copula density, i.e. a PDF of a distribution with uniform marginals corresponding to the CDF *C*(*u, v*). Hereafter, we will use “copula” to refer to *C*(*u, v*) and “copula density” to refer to its derivative, 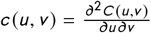.

To draw samples from a copula, which can then be used to draw samples from a joint distribution with prespecified marginals, we follow a procedure based on conditional distributions (Johnson, 2013). The conditional distribution of a copula is defined as

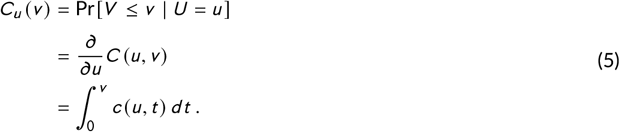

To illustrate, we will draw samples from a Gaussian copula and a Clayton copula (left and right columns of Fig. 2, respectively). A definition of the Clayton copula, together with other copulae of the Archimedean family can be found in the appendix, section A2. To generate a sample from *C*(*u, v*), we first draw *u* from a uniform distribution and then draw *v* from *C_u_*(*v*). The pair (*u, v*) is then a sample from *C*(*u, v*). To draw *v* from *C_u_*(*v*), we draw *z* from a uniform distribution and use the probability integral transform to obtain 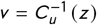. We repeat this process *n* times to generate a sample of size *n* from *C*(*u, v*) (Fig. 2 a,b). We can then transform this sample (*u, v*) from the copula to a sample from a bivariate distribution, (*x, y*), with non-uniform marginal CDFs *F_X_*(*x*) and *F_Y_*(*y*) via 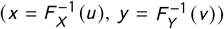 (Fig. 2, second row, for normally distributed margins and third row for gamma and exponentially distributed margins).

Finally, any copula is bounded by the Fréchet-Hoeffding bounds (Fréchet, 1935; Hoeffding, 1940; Fréchet, 1951)

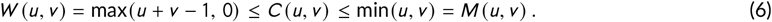

*W* and *M* are themselves copulae (at least in the bivariate case). However, some copulae do not attain one or either bound, in which case they are not capable of capturing strong positive or negative correlations. The Gaussian copula, for example, approaches the lower and upper Fréchet-Hoeffding bounds for *ρ* → −1 and *ρ* → 1, respectively, and is equal to the product copula for *ρ* = 0 (see appendix, Figure A1); the product copula Π (*u, v*) = *uv* is a copula corresponding to independent random variables. The Clayton copula, on the other hand, cannot reach the lower Fréchet-Hoeffding bound, approaches the upper bound as *α* → ∞, and approaches the product copula as *a* → 0 (from above or below). For those interested in learning more about copulae, we recommend the following excellent reviews (in our subjective order of increasing mathematical difficulty): Trivedi and Zimmer (2007), Nelsen (2006), Joe (2014); for readers with a working knowledge of measure theory, we recommend Durante and Sempi (2015).

### 3.1 Circular-Linear Copulae

Our main objective is to develop appropriate copulae for modeling joint turn-angle and step-length distributions. Circular-circular copulae and circular-linear copulae without the restriction of symmetry (see below) have been a subject of statistical literature starting with Johnson and Wehrly (1977) and Johnson and Wehrly (1978), who, however, did not yet use the term “copula”. With these copulae still being an active area of research, we will also draw inspiration from more recent reviews (e.g. Jones et al., 2015) and applied studies (e.g. García-Portugués et al., 2013). Let Θ be a continuous circular random variable and *X* a continuous linear one. While *X* has support on the real line, 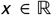, Θ has support on the unit circle, 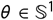, which we will view as having support on some interval of length 2*π*, [*a, a* + 2*π*). The PDF then needs to fulfill the boundary condition

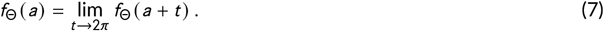

This condition is somewhat different from (though equivalent to) the definitions found in standard references, such as Mardia and Jupp (2000), where they view 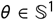 as having support on the entire real line, which means that the PDF needs to be 2*π*-periodic. The first definition is more convenient for our purposes, since it mirrors the definition of the density of a circular-linear copula (see equation 8). Examples of circular distributions commonly used in movement studies include von Mises and wrapped Cauchy distributions (see Watson, 1983; Mardia and Jupp, 2000; McClintock et al., 2012).

Any continuous bivariate circular-linear PDF has support on (a subset of) the cylinder, 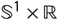. From equation 4 it is evident that this means that also the domain of the corresponding copula is not the unit square, but the surface of a cylinder of unit height and unit circumference and

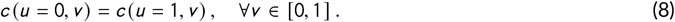

For the remainder of the text, we will define *u* to be the circular and *v* the linear dimension of the copula density, and for the sake of brevity, we will refer to copulae with densities fulfilling equation 8 as “periodic in u”, or just as “periodic”.

We will set the support of the turn angle Θ to [-*π, π*), with negative angles corresponding to left turns and positive angles to right turns. The support of the linear random variable *X*, the step length, is restricted to the non-negative real numbers. Turn angles are somewhat different from other circular variables in that it often makes sense to assume that an animal has no inherent bias to turn left or right. This can be achieved with a marginal PDF that is symmetric around *θ* = 0 and a copula that has not only a periodic density (see equation 8), but also a symmetric one in direction *u* around *u* = 0.5

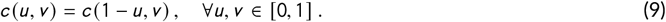

For the sake of brevity, we will refer to copulae having the property in equation 9 as “symmetric in u”, or just as “symmetric”, even though there are many other possible symmetries for copulae (see Nelsen 2006 and Hodel and Fieberg 2021). Finally, note that any copula that is “symmetric” will also be “periodic”.

## 4 CONSTRUCTING CIRCULAR-LINEAR COPULAE

Johnson and Wehrly (1978) showed that circular-linear densities with given marginals can be obtained via

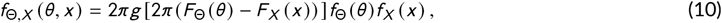

where *g*(*ω*) is a 2*π*-periodic density function on the circle. Thus, many have recognized that *c*(*u, v*) = 2*πg* [2*π* (*u – v*)] is a circular-linear copula density (see e.g. García-Portugués et al., 2013). We have implemented this copula using a von Mises density for *g*(*ω*) (see `cyl_vonmises` in our `cylcop` package (Hodel and Fieberg, 2021)). However, there is no obvious way to obtain symmetric joint densities using this approach, which makes it less useful for movement data. We will therefore introduce in the following sections other methods of obtaining copulae with densities not only periodic, but also symmetric in *u*.

### 4.1 Copulae with Quadratic or Cubic Sections

The section in *v* at *u*_0_ of a copula *C*(*u, v*) is a curve *s*_*u*_0__(*v*) = *C*(*u*_0_, *v*) that is defined by the vertical cross-section of the copula where *u* is held constant at *u* = *u*_0_. Consider the following class of circular-linear copulae, for which the sections in the linear dimension *v* are quadratic functions

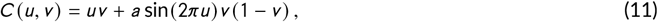

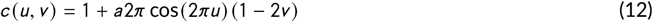

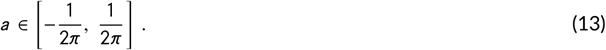

It is easy to see that *c*(*u, v*) is not only periodic in *u*, but also symmetric

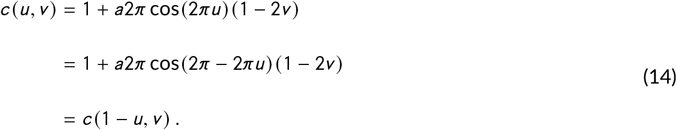

Similarly, the class of circular-linear copulae, below, with cubic sections in the linear dimension *v* will also be periodic and symmetric in the circular direction *u*

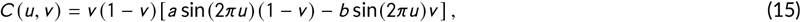

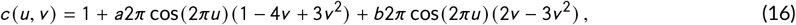

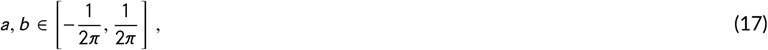

since cos(2*πu*) = cos(2*π*(1 – *u*)). Copulae with quadratic and cubic sections are implemented in the `cylcop` package as `cyl_quadsec` and `cyl_cubsec` and further details including the derivation of their densities and conditional distribution functions are given in Hodel and Fieberg 2021. An example of a copula with quadratic sections in *v* is displayed in figure 3 and an example of one with cubic sections is shown in Figure 6a. To fully appreciate the difference between copulae with quadratic and cubic sections, we refer the reader to our app as well as Hodel and Fieberg 2021.

**FIGURE 3.**
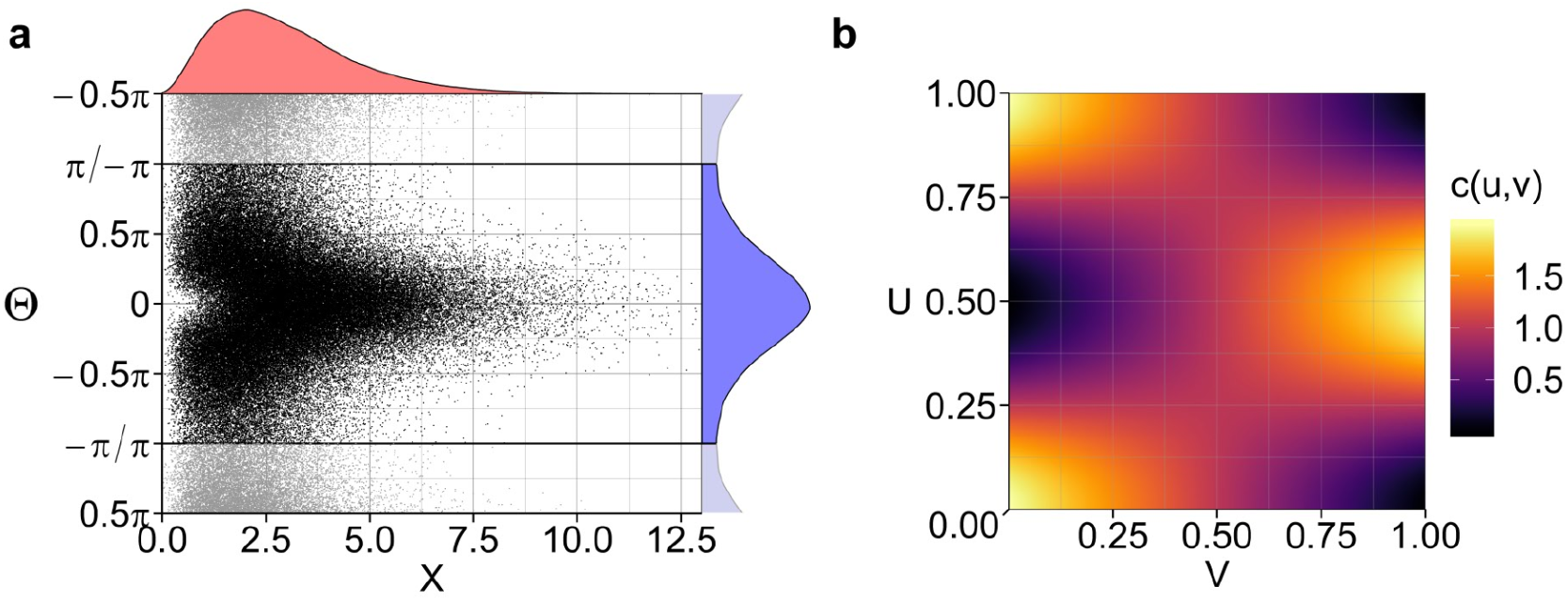
Left: linear (*x*) and circular (*θ*) samples drawn from a joint distribution obtained using a copula with quadratic sections in *v* (equation 12, with *a* = 1 /(2*π*)), a marginal gamma distribution (shape=3, scale=1), and a marginal von Mises distribution (*μ* = 0, *κ* = 1). Right: PDF of the quadratic-sections copula with *a* = 1 /(2*π*).

### 4.2 Combinations of Copulae

A second way to generate circular-linear copulae that are symmetric in *u* is to combine two or more linear-linear copulae. In doing so, we will make use of the following properties of copulae (Nelsen, 2006):

1. Any convex linear combination of copulae is a copula.
2. Orthogonal reflections of a copula density *c*(*u, v*) with respect to the lines *u* = 0.5 or *v* = 0.5 are again densities of a copula.

#### 4.2.1 Linear Combinations of Reflected Copulae

In some special cases, orthogonal reflections can also be seen as rotations by a multiple of 0.5*π* (see Hodel and Fieberg 2021). It is therefore common (see e.g. Patton, 2012; Yoshiba, 2016; Hofert et al., 2018) to denote a copula obtained by reflecting the density of copula *C* along the line *u* = 0.5 by *C*_0.5*π*_, along the line *v* = 0.5 by *C*_1.5*π*_, and along both lines *u* = 0.5 and *v* = 0.5 by *C_π_*; these copulae are often referred to as “rotated copulae”.

The density and distribution of the copula generated by reflecting *c*(*u, v*) with respect to *u* = 0.5 are

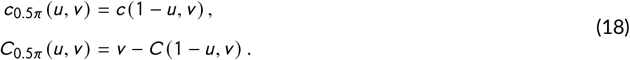

The CDF can be derived easily by integrating the PDF and taking care of the boundary conditionsofa copula (see Hodel and Fieberg 2021). Taking the arithmetic mean between any linear-linear copula *C* and *C*_0.5*π*_ produces a circular-linear copula

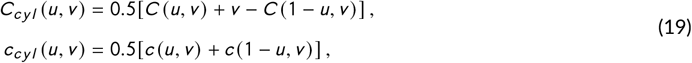

with a density that is not only periodic, but also symmetric in *u*. This density, which usually has an X-shaped appearance, can also be “periodically shifted” by 0.5 in the *u*-direction, which gives a copula with diamond-shaped density that is also symmetric and periodic (appendix, Fig. A5). The structure of the correlation captured by such copulae in the context of movement data is as follows: Small values of step lengths correlate with turn angles around *π* or, equivalently, −*π*, step lengths close to the median (i.e. step lengths close to relative rank 0.5) correspond to turn angles close to 0, and large step lengths correlate again with large turn angles around *π* or −*π*. For the shifted, “diamond-shaped” copulae the opposite is true, with small and large step lengths associated with small turn angles. This approach is implemented in the `cylcop` package as `cyl_rot_combine` (Hodel and Fieberg, 2021). For further illustrations, we refer the reader to our app (https://cylcop.shinyapps.io/cylcop-graphs/).

#### 4.2.2 Rectangular Patchworks of Copulae

The last method for generating symmetric circular-linear copulae entails partitioning the unit square into rectangular regions, each containing an appropriately transformed copula. To construct such rectangular patchworks of copulae, we follow the procedure outlined in Durante et al. (2009). Specifically, we define 2 rectangles that are symmetric about *u* = 0.5: *R*_1_ = [*u*_1_, *u*_2_] × [0,1] and *R*_2_ = [1 – *u*_2_, 1 – *u*_1_] × [0, 1] with 0 ≤ *u*_1_ < *u*_2_ ≤ 0.5. Next, let *C_bg_*(*u, v*) (“bg” for background) be a copula and *F_i_*(*u, v*): *R*, → [0, 1] a 2-increasing function obtained by some transformation of a copula *C_i_*(*u, v*) so that *F_i_* = *C_bg_* at the boundaries of each rectangle, *R_i_*. The patchwork-copula is then

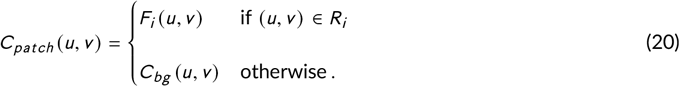

To generate a copula *C_patch_* that is periodic and symmetric in *u*, the two copulae from which the functions *F_i_* are derived must be reflections of each other with respect to the line *u* = 0.5. This can be achieved by defining *C*_2_ (*u, v*) = *C*_1,0.5*π*_(*u, v*) = *v* – *C*_1_(1 – *u, v*). With this condition satisfied, we then either set *u*_1_ = 0, *u*_2_ = 0.5 (see upper row of Fig. 4), or we can choose a copula that is periodic and symmetric in *u* for *C_bg_* and use any values of *u*_1_ and *u*_2_ that we desire (see lower row of Fig. 4). For illustrative purposes, we have chosen copulae with quite extreme parameter values. For a rectangular patchwork copula with smoother density, which is usually more appropriate for movement data, see Figure A9f in the appendix. The explicit distribution of a general patchwork-copula (of which the above-described copula is a special case) is given in theorem 2.2 in Durante et al. (2009). The equations and derivations for our two-rectangle-copulae can be found in Hodel and Fieberg 2021, and they are implemented in the `cylcop` package as `cyl_rect_combine`.

**FIGURE 4.**
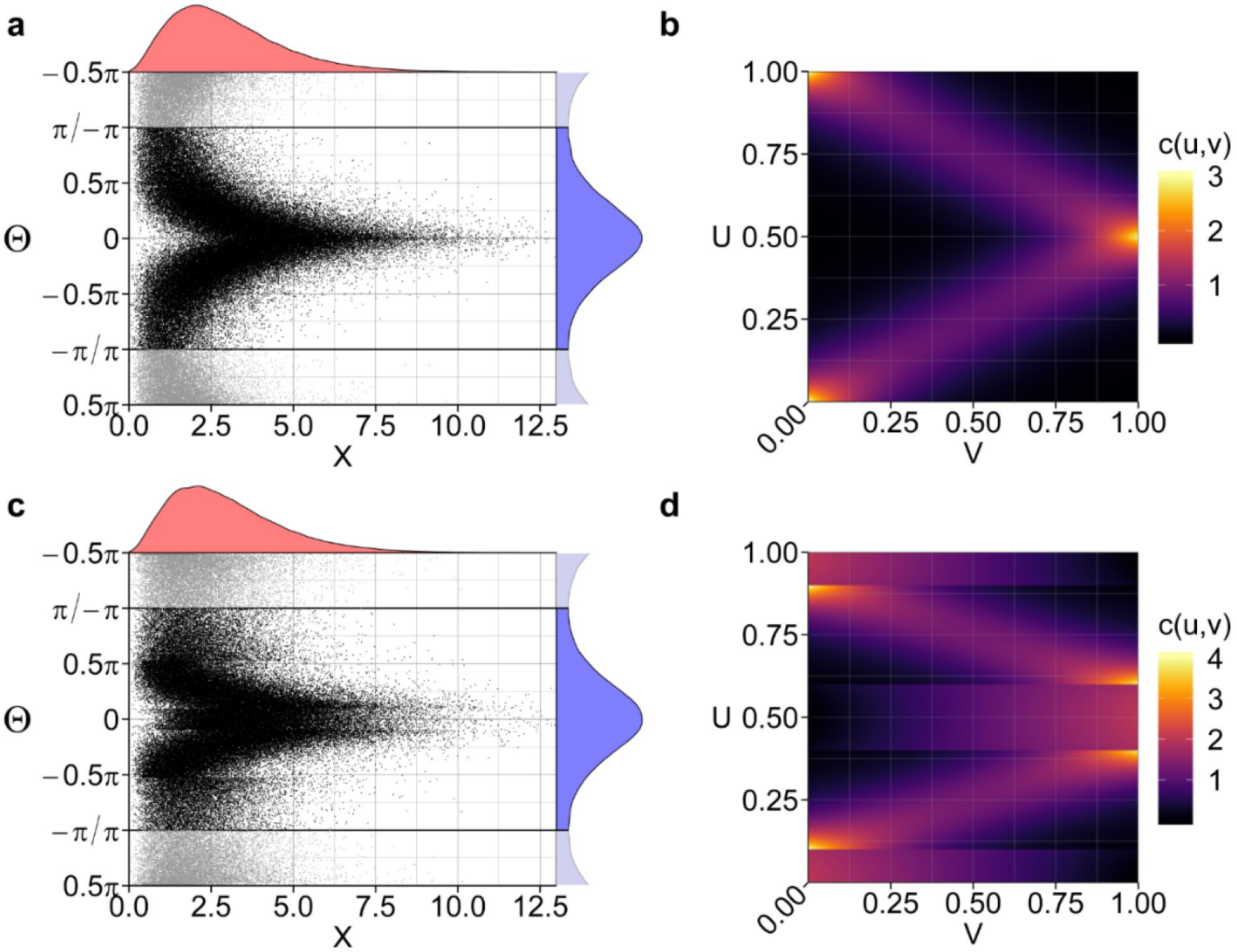
Left column: step lengths (*x*) and turn angles (*θ*) drawn from joint distributions obtained with `cyl_rect_combine`-copulae, a marginal gamma distribution (shape=3, scale=1), and a von Mises distribution (*μ* = 0, *κ* = 1). Right column: PDFs of `cyl_rect_combine`-copulae. Upper row: The rectangular patchwork of the copula consists of the 2 rectangles: *R*_1_ = [0, 0.5] × [0, 1] and *R*_2_ = [0.5, 1] × [0, 1]. The function in the lower rectangle is obtained by a transformation of a Frank copula (Frank, 1979) with *α* = 8 and the function in the upper rectangle by transforming a 90 degrees rotated Frank copula with *α* = 8. Lower row: The rectangular patchwork of the copula consists of the 2 rectangles: *R*_1_ = [0.1, 0.4] × [0, 1] and *R*_2_ = [0.6, 0.9] × [0, 1]. The function in the lower rectangle is obtained by a transformation of a Frank copula (*α* = 8) and the function in the upper rectangle by transforming a 90 degrees rotated Frank copula with *α* = 8. The “background copula” is a `cyl_quadsec`-copula with parameter *a* = 1 / (2*π*).

## 5 PARAMETER ESTIMATION AND DEPENDENCE

Although it is possible, in principle, to estimate parameters of the two assumed marginal distributions and the copula simultaneously by maximizing the log-likelihood using the joint PDF of equation 4, this leads to a complex optimization problem. Further, estimators of copula parameters can be sensitive to mis-specification of the marginal distributions, and estimators of parameters in the marginal distribution can be influenced by mis-specification of the copulae. Instead, we implement a maximum pseudo-likelihood estimator (MPLE, see Oakes, 1994; Genest et al., 1995; Shih and Louis, 1995; Tsukahara, 2005). We begin by finding the optimal parameters of the marginal distributions via maximum likelihood estimation. Next, we calculate a non-parametric estimate for each marginal distribution and then use the resulting empirical marginal CDFs, 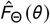 and 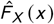, to transform the set of *n* i.i.d. observations of step-lengths and turn angles (*θ_i_, x_i_*) to the pseudo-observations 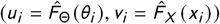, i.e. values drawn from the empirical copula. Finally, we use MPLE to fit the parameters *α* of our selected copula family to the pseudo-observations by maximizing

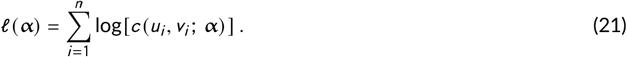

Compared to maximum likelihood estimations using the full joint distribution, this MPLE approach reduces the number of parameters that have to be optimized simultaneously. It is important to provide “good” starting parameter values for MPLE, especially for rugged parameter hyper-surfaces, and we derive them by comparing circular-linear correlation measures for the empirical and parametric copula with different parameter values (see Hodel and Fieberg 2021 for further details). For rectangular patchwork copulae with rectangles spanning the entire unit square, parameter values (i.e. starting values for MPLE) can be derived analytically from Kendall’s tau of the empirical copula (Oakes, 1982; Genest, 1987; Genest and Rivest, 1993). Finally, model selection can be performed using Akaike’s information criterion, which is a common (e.g. Chen and Fan, 2005; McNeil et al., 2015; Joe, 2014), but for MPLE not uncontroversial approach (see Grønneberg and Hjort 2014, but also Jordanger and Tjøstheim 2014).

### 5.1 Direction of Correlation

For all symmetric copulae introduced in this paper, periodically shifting the density by 0.5 in *u*-direction leads again to a symmetric copula. This “shifting” is implemented differently in all copulae: for copulae with quadratic or cubic sections, it means multiplying their parameters by −1. For the patchwork-copulae, it means reflecting or rotating by 90 degrees *c*_1_ and *c*_2_ (and adapting the background copula, if applicable). Finally, for the copulae obtained by taking the arithmetic mean of a linear-linear copula and its reflected counterpart, we implemented the “shifting” explicitly to go from an x-shaped to a diamond-shaped density, as mentioned in section 4.2.1. In any case, periodically shifting the density reverses the correlation between the two random variables. If a copula captures e.g. a correlation of large step lengths with small absolute angles, the shifted copula corresponds to a correlation of small step lengths with small absolute angles.

## 6 APPLICATION TO FISHER LOCATION DATA

### 6.1 Methods

We fitted bivariate distributions to the fisher data displayed in Figure 1 using the copulae and methods introduced in the previous sections. We considered several different linear and circular marginal distributions (see appendix, tables A1 and A2) and selected the appropriate ones using maximum likelihood. Next, we considered several different copuale based on the characteristics of the data. With our assumption that an animal has no inherent bias to turn left or right, we can ignore the (only briefly mentioned) circular-linear copula by Johnson and Wehrly, since it is not symmetric in *u*. A visual inspection of the data further excludes copulae obtained from linear combinations of reflected copulae with their x- or diamond-shaped densities. We therefore decided to fit copulae with quadratic (`quad_sec`) and cubic sections (`cub_sec`), as well as 3 rectangular patchwork copulae. The latter were restricted to consist of two symmetric rectangles, together covering the entire unit square. They were based on Clayton (`rect_combine_Clayton`), Gumbel (`rect_combine_Gumbel`) and Frank copulae (`rect_combine_Frank`), three commonly used Archimedean copulae (see appendix, section A2 for a quick overview of their properties). The functions in the lower rectangles were derived directly from those copulae, the ones in the upper rectangles from their 90 degree rotations. We also attempted to fit a rectangular patchwork copula based on the Frank copula, again with two symmetric rectangles but this time with their lower and upper bound as tuneable parameters and a quadratic sections copula as the background copula.

We generated preliminary estimates of copula parameters using measures of circular-linear correlation (see Hodel and Fieberg 2021), which we then used as starting values for MPLEs. We determined starting values for `quad_sec` and `cub_sec` using a grid-search, choosing parameter values that gave the best match to the circular-linear correlation coefficient of the data; starting values for the `rect_combine`-copulae were chosen to match Kendall’s tau of the underlying linear-linear copulae to the data and could be calculated analytically. When fitting the rectangular patchwork copula with lower and upper bounds as tuneable parameters, the MPLE converged either to `rect_combine_Frank` or `quad_sec` depending on the starting values. Therefore, purely for illustrative purposes, we fixed the rectangles to *R*_1_ = [0.0,0.3] ×[0,1] and *R*_2_ = [0.7,1.0] ×[0,1] and optimized the parameters of the Frank and the quadratic sections copula (`rect_combine_Frank-quad_sec`). We compared copulae using AIC and visually investigated their properties and the properties of the corresponding joint distributions using various plots. We also generated trajectories of different lengths and estimated diffusion coefficients, *D*, by regressing mean-square displacement against time for the `cub_sec` copula and for simulations using independent samples of turn angles and step lengths (appendix, Section A4.2).

### 6.2 Results

The marginal distribution of the step lengths was best described by a log-normal distribution with parameters *μ* = 3.83 and *σ* = 1.41. As already remarked in the introduction, the marginal distribution of the turn angles is bimodal and the smallest AIC score was achieved with a mixture of two von Mises distributions. We fixed the location parameters at 0 and *π* and estimated the concentration parameters to be 0.48 and 6.31, respectively, with a mixing proportion of 0.78 (see appendix, tables A1 and A2).

Of the copulae we considered, the cubic sections copula (`cub_sec`) had the lowest AIC score (Table 1). To visualize it, we simulated 4350 step-lengths and turn angles from the `cub_sec` copula and from the independence copula (Figures 5 and 6; data simulated with the other copulae can be found in the appendix, Figure A7). Visually, it is clear that the data generated with `cub_sec` (Figure 5a) provide a better match to the “real” fisher data (Figure 1a) than the data simulated with independent turn angles and step lengths (Figure 5c). The effect of the correlation structure can also be seen from the circular box-plots of the turn angles (Figures 1b, 5b and 5d). We calculated the pseudo-observations of the fisher-data and smoothed them using a two-dimensional kernel density estimator (KDE) with a bivariate Gaussian kernel. While this KDE is a helpful visualization, note that it is mathematically not a valid empirical, non-parametric (circular-linear) copula (see Section 7). The KDE and the density of `cub_sec` both highlight that large step lengths tend to be associated with turn angles near 0 and small step lengths with turn angles near ±*π* (Fig. 6). Plots of copula densities for the other copulae (Table 1) can be found in the appendix (Figure A9). The `cub_sec` copula resulted in a diffusion coefficient, *D* = 12.8 m^2^/s, significantly smaller than the diffusion coefficient assuming independent steps and turns, *D* = 17.7 m^2^/s.

**FIGURE 5.**
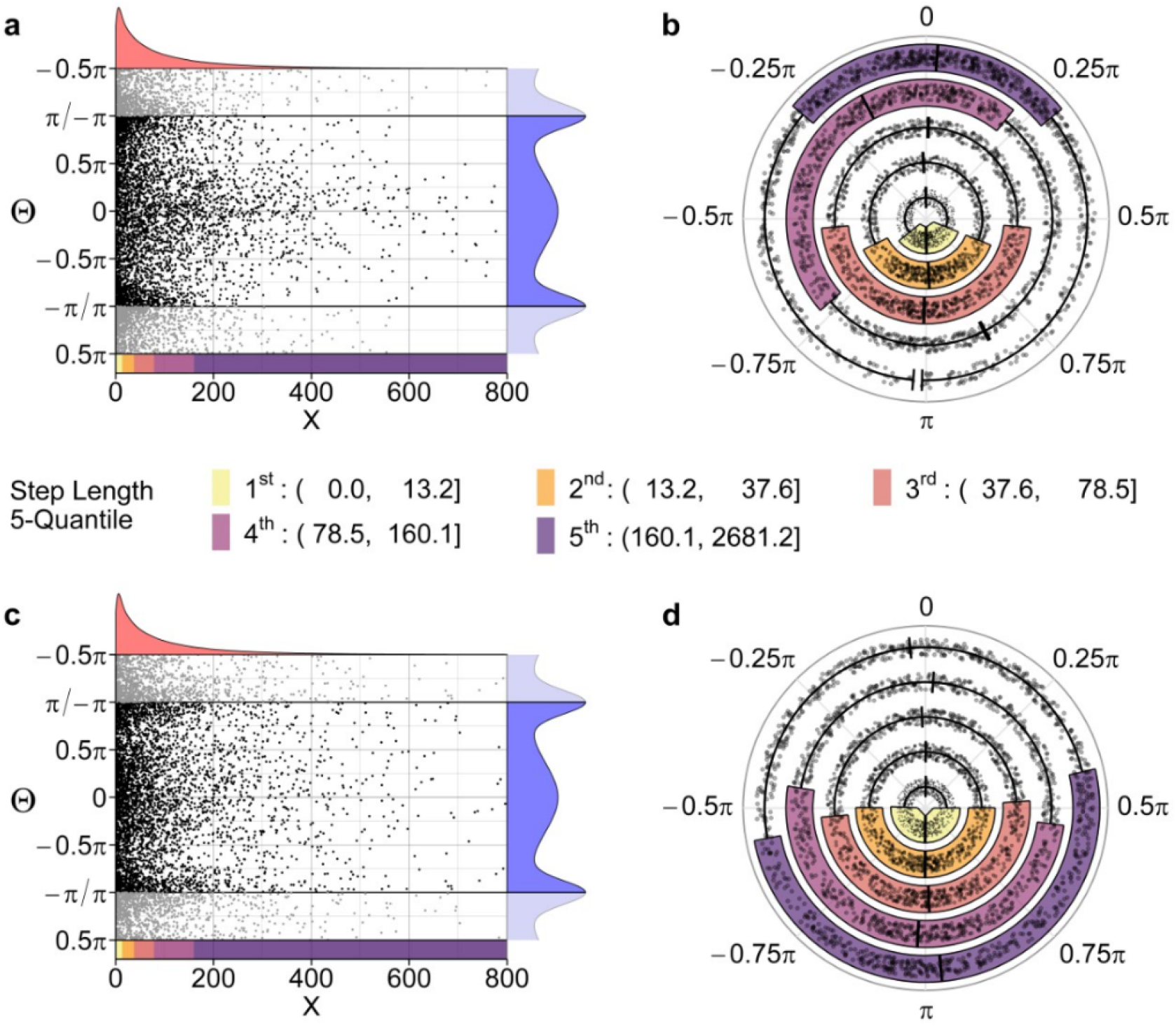
Top row: 4350 step lengths and turn angles sampled from a joint distribution obtained with the cubic sections copula `cub_sec`. Bottom row: 4350 step lengths and turn angles sampled independently from their marginal distributions. Left column: Scatter plots of step lengths and turn angles. Maximum likelihood estimates of the marginal densities are plotted in red and blue next to the axes. The step lengths are separated into 5 quantiles as marked by the colorbar below the X-axis. Right column: Circular box plots of the same data. For each of the 5 step length quantiles, a box plot of the corresponding angles is shown.

**TABLE 1.**
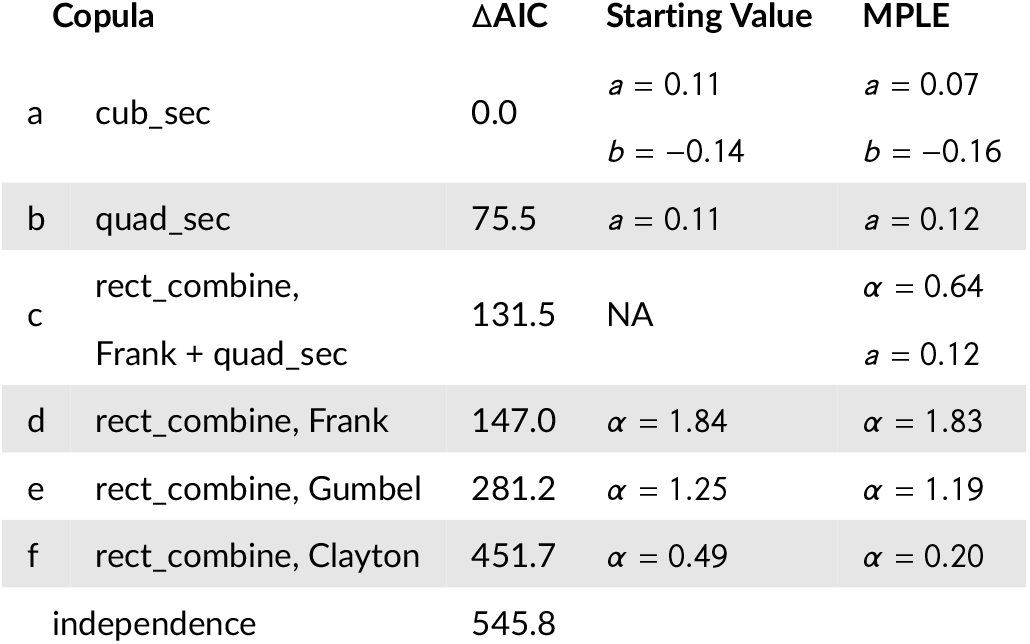
AIC relative to the lowest AIC (`cub_sec`) and maximum pseudo-likelihood parameter estimates for 6 copulae fit to the fisher data. Starting values for copulae a and b were determined using a grid-search, choosing parameter values that gave the best match to the circular-linear correlation coefficient of the data; starting values for copulae d-f were chosen to match Kendall’s tau of the data.

| Copula | $\Delta AIC$ | Starting Value | MPLE |
| --- | --- | --- | --- |
| a *cub_sec* | 0.0 | $a = 0.11$<br>$b = -0.14$ | $a = 0.07$<br>$b = -0.16$ |
| b *quad_sec* | 75.5 | $a = 0.11$ | $a = 0.12$ |
| c *rect_combine*,<br>*Frank + quad_sec* | 131.5 | NA | $\alpha = 0.64$<br>$a = 0.12$ |
| d *rect_combine*, *Frank* | 147.0 | $\alpha = 1.84$ | $\alpha = 1.83$ |
| e *rect_combine*, *Gumbel* | 281.2 | $\alpha = 1.25$ | $\alpha = 1.19$ |
| f *rect_combine*, *Clayton* | 451.7 | $\alpha = 0.49$ | $\alpha = 0.20$ |
| independence | 545.8 |  |  |

The estimate of the parameter *b* of the `cub_sec` copula was on the boundary, and its circular-linear correlation coefficient was slightly lower than the coefficient for the actual fisher data (0.085 versus 0.102). These results suggest that a closer fit might be possible using a copula with a correlation structure similar to the `cub_sec` copula, but with a wider range of possible dependencies. The copulae consisting of rectangular patchworks can potentially capture any degree of correlation between both Fréchet–Hoeffding bounds and a wide range of joint density shapes. Thus, it may be beneficial to consider other correlation structures beyond the three Archimedean copulae we selected. The main drawback of these type of copulae, however, is that they can be difficult to optimize since one has to choose and compare copulae for the patches and the background, and estimate parameters for each of these copulae. Introducing more than two rectangular patches is straight forward but would further complicate optimization.

## 7 DISCUSSION AND FUTURE RESEARCH

The observed correlation between step-lengths and turn angles in our applied example is typical of many modern-day telemetry data sets. We have shown that this correlation can be captured by new circular-linear copulae that we have developed to also mimic characteristics of animal movement data (e.g., symmetric turn angle distributions). We have demonstrated methods for estimating copulae parameters and for simulating data from fitted models. Lastly, we have shown that this correlation can impact movement patterns (e.g., dispersion over time).

We see several opportunities to expand on our work, including opportunities for both theoretical and methodological advances as well as research that will allow copulae to be incorporated into existing frameworks for modeling animal movement and habitat use. One of the main characteristics of movement data, which we have so far neglected, is temporal autocorrelation. Although turn-angle distributions with modes at 0 allow *steps* to be autocorrelated (i.e., they allow individuals to maintain a consistent direction), the copulae we have considered do not allow turn angles or step-lengths themselves to be autocorrelated. To incorporate this feature, our MPLE procedure would need adapting to allow for non-i.i.d. data. Often, covariates are recorded along with movement metrics, and the target of modeling is the joint distribution of step lengths and turn angles conditional on the covariates, or, in other words, regression. Similarly, in time series analysis one is often interested in the distribution of a random vector conditional on past instances. Both issues can be tackled using conditional copulae (Patton, 2006; Fermanian and Wegkamp, 2012).

**FIGURE 6.**
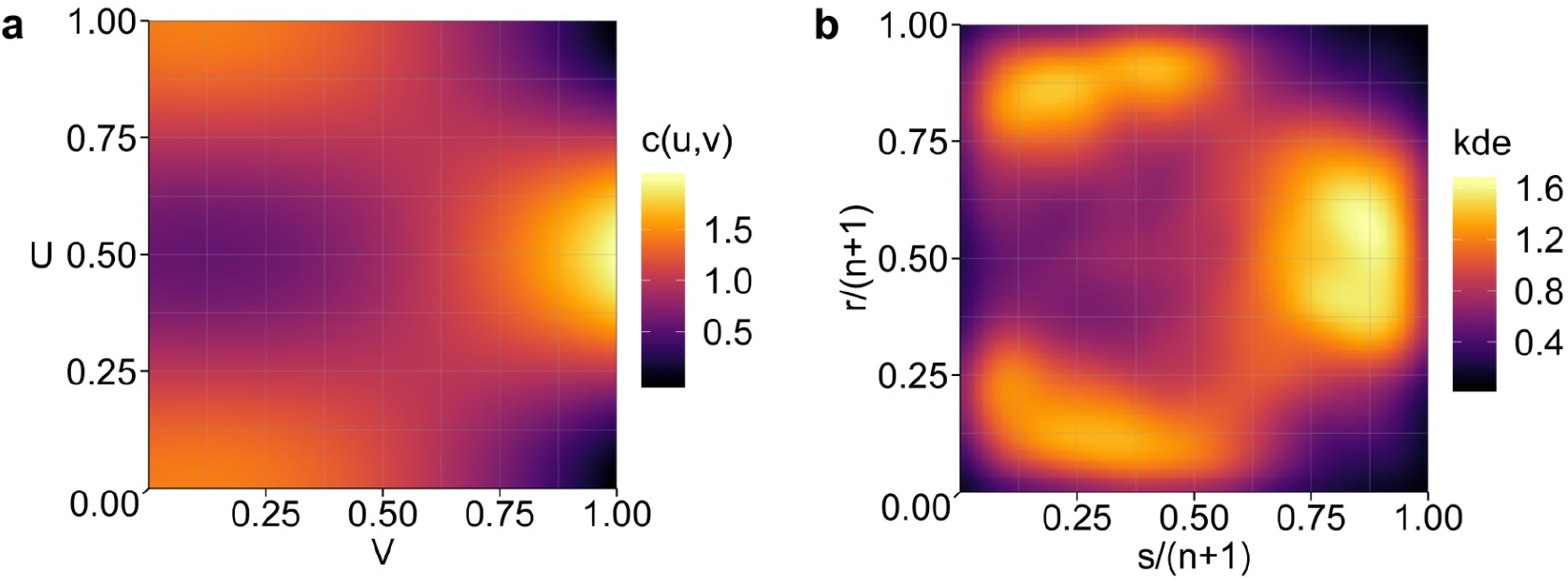
Left: Density of the cubic sections copula `cub_sec`. Right: Kernel density estimate of the pseudo-observations of the fisher data.

We have focused this paper on parametric copulae with parameters estimated using frequentist methods. Distributions of copula parameters could also be estimated in a fully Bayesian framework. Alternatively, copulae can be estimated non-parametrically by smoothing the empirical copula using e.g. Bernstein polynomials (Sancetta and Satchell, 2004; Janssen et al., 2012). This has already been done in the general circular-linear case (see Carnicero et al. 2013; García-Portugués et al. 2013, 2014) and would need only slight adaptation for our symmetric circular-linear data.

Other areas where additional research might be rewarding include the development of graphical tools to assess goodness-of-fit or to visualize dependence, such as Chi-plots (Fisher and Switzer, 1985, 2001) or K-plots (Genest and Boies, 2003). These could be implemented along with a cross-validation procedure for assessing goodness-of-fit (Grønneberg and Hjort, 2014). A more thorough investigation of correlation measures applicable to movement data and the possibility of estimating copula parameters from them could help in pre-selecting copula families and provide better initial guesses for MPLE. Lastly, we see many opportunities for integrating these methods into existing frameworks for modeling animal movement. In particular, copula could be used to more accurately sample available steps when conducting step-selection analyses (Fortin et al., 2005; Forester et al., 2009; Thurfjell et al., 2014; Fieberg et al., 2021) or to better capture movement modes in hidden Markov or state space models (Morales et al., 2004; Patterson et al., 2008; McClintock and Michelot, 2018; McClintock et al., 2020).

## Supporting information

Code application

## Acknowledgements

JF received partial salary support from the Minnesota Agricultural Experimental Station and from the Minnesota Environmental and Natural Resources Trust Fund as recommended by the Legislative-Citizen Commission on Minnesota Resources (LCCMR). We thank Dr. Gabriele Cozzi for his valuable input.

## Conflict of Interest

None.

## Data Availability

Upon acceptance, data and code will be archived in the Data Repository of the University of Minnesota (https://conservancy.umn.edu/handle/11299/166578).

## Notes

https://github.com/floo66/cylcop

https://cylcop.shinyapps.io/cylcop-graphs/

### Competing Interest Statement

The authors have declared no competing interest.

