## Appendix for "Circular-Linear Copulae for Animal Movement Data"

#### A1 Copula

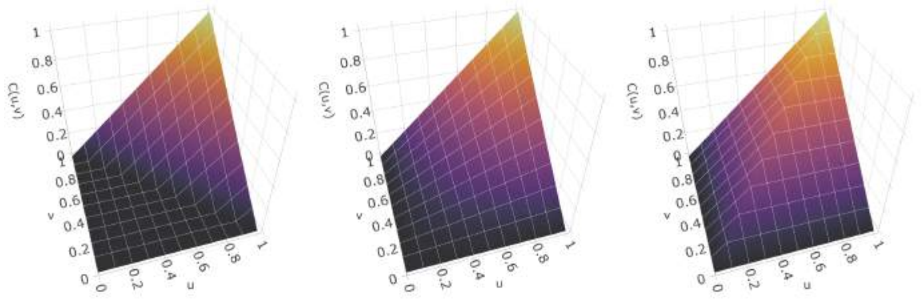


Figure A1: Surface plots of the lower Fréchet–Hoeffding bound $W(u,v)$ (left), the product copula $\Pi(u,v)$, and the upper Fréchet–Hoeffding bound $M(u,v)$ (right).

#### A2 Archimedean Copulae

Following chapter 4 in (Nelsen 2006), Archimedean copulae can be generated according to

$$C(u,v) = \varphi^{[-1]}(\varphi(u) + \varphi(v))\,, \tag{1}$$

where $\varphi : [0,1]^2 \to [0,\infty]$ must be a continuous, strictly decreasing, convex function, such that $\varphi(1) = 0$ with it's pseudo-inverse $\varphi^{[-1]}$ defined as

$$\varphi^{[-1]}(t) = \begin{cases} \varphi^{-1}(t) & \text{if } t \in [0, \varphi(0)] \\ \varphi(0) & \text{if } t \in [\varphi(0), \infty]\,. \end{cases} \tag{2}$$

The three most commonly used Archimedean copulae are the Frank, Clayton and Gumbel copula.

#### A2.1 Frank Copula

This symmetric copula is obtained by setting

$$\varphi_\alpha(t) = -\ln\left(\frac{\exp(-\alpha t) - 1}{\exp(-\alpha) - 1}\right) \qquad \text{with } \alpha \in (-\infty, \infty) \setminus \{0\} \tag{3}$$

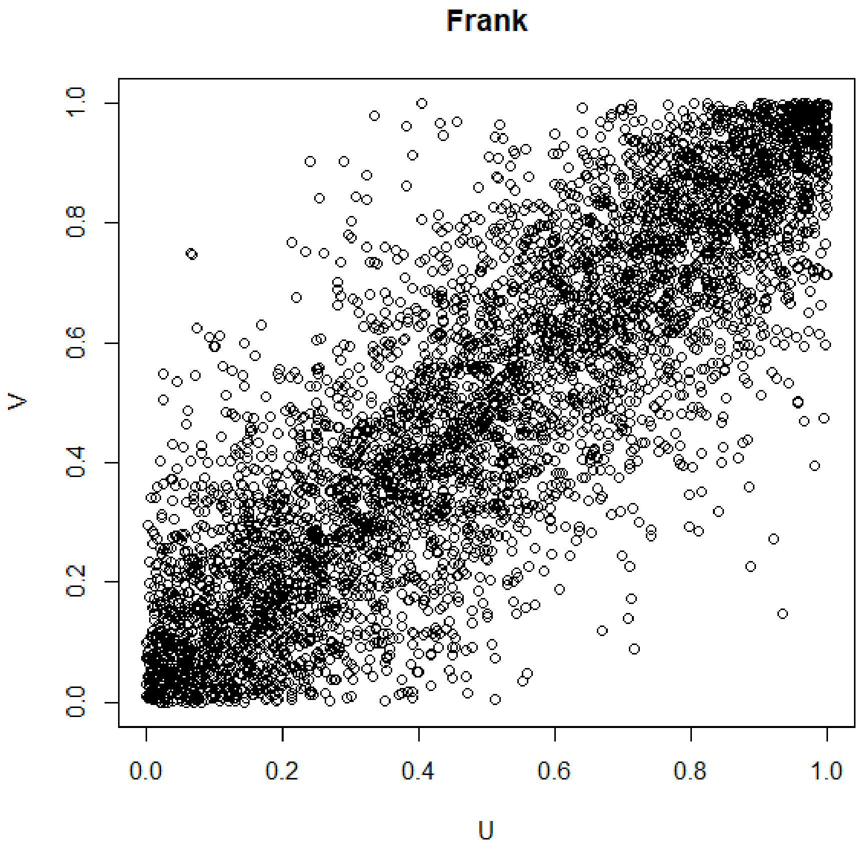


Figure A2: Frank copula with $\alpha = 8$

#### A2.2 Clayton Copula

This asymmetric copula with greater dependence in the negative tail (small values of $u$ and $v$) than in the positive tail is obtained by setting

$$\varphi_\alpha(t) = \frac{1}{\alpha}\left(t^{-\alpha} - 1\right) \qquad \text{with } \alpha \in (-1, \infty) \setminus \{0\} \tag{4}$$

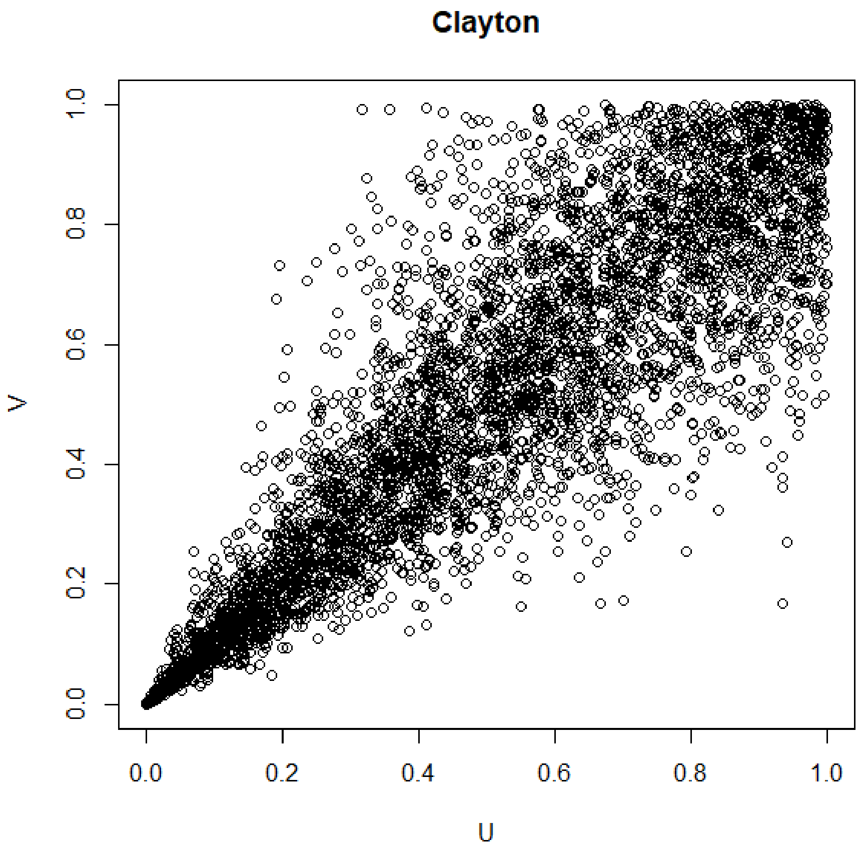


Figure A3: Clayton copula with $\alpha = 5$

##### A2.3 Gumbel Copula

This asymmetric copula with greater dependence in the positive tail (large values of $u$ and $v$) than in the negative tail is obtained by setting

$$\varphi_\alpha(t) = (-\ln(t))^\alpha \qquad \text{with } \alpha \in (1, \infty) \tag{5}$$

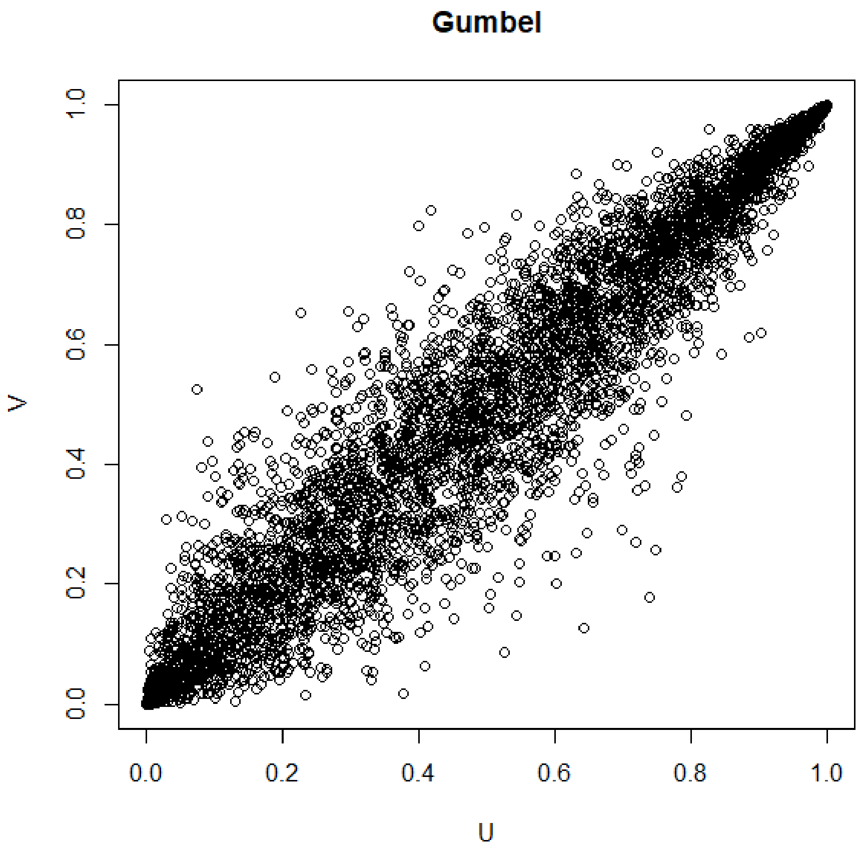


Figure A4: Gumbel copula with $\alpha = 5$

#### A3 Linear Combination of Reflected Copulae

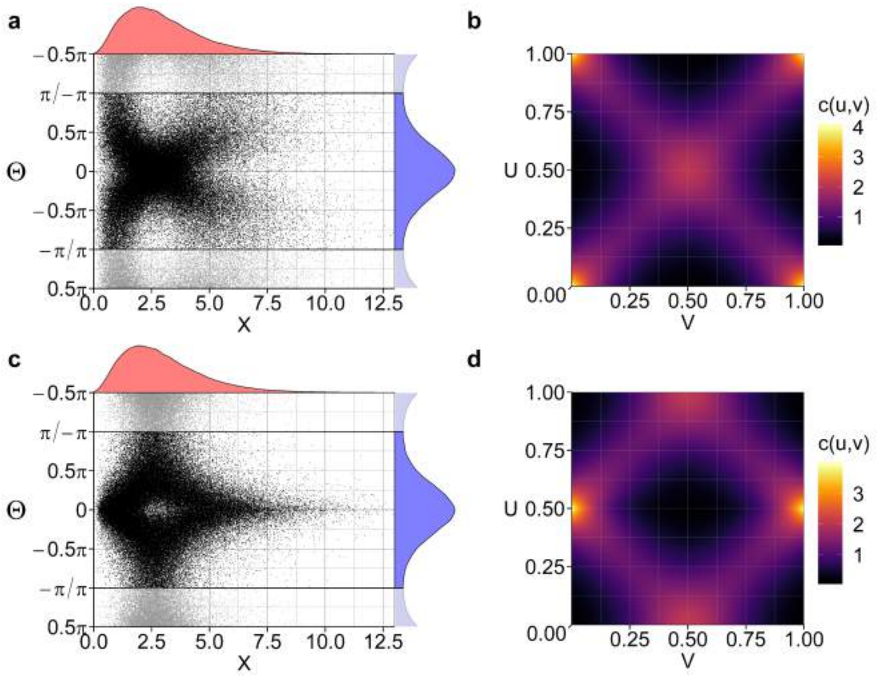


Figure A5: a: linear ($x$) and circular ($\theta$) samples drawn from a joint distribution obtained with a `cyl_rot_combine`-copula construced as the arithmetic mean of a Frank copula and a 90 degrees rotated Frank copula, both with $\alpha = 8$. The marginals follow a gamma distribution (shape=3), and a von Mises distribution ($\mu = 0$, $\kappa = 1$). b: PDF of that `cyl_rot_combine`-copula. c: samples drawn from a joint distribution obtained with the same marginals and copula parameters, but with the copula density shifted periodically by 0.5 in u-direction. d: PDF of that copula.

### A4 Application to Fisher Data

Table A1: AIC and maximum likelihood estimates of parameter values from fitting circular distributions to the turn angles of the fisher data.

| **Distribution** | **AIC** | **Parameters** | |
|---|---|---|---|
| von Mises | 15991 | $\mu = -2.86$ | $\kappa = 0.05$ |
| wrapped Cauchy | 15987 | $\mu = 3.33$ | $\rho = 0.03$ |
| mixed von Mises | 15593 | $\mu_1 = -0.05$<br>$\kappa_1 = 0.48$<br>prop $= 0.78$ | $\mu_2 = 3.13$<br>$\kappa_2 = 6.38$ |
| mixed von Mises | 15590 | $\mu_1 = 0$ (fixed)<br>$\kappa_1 = 0.48$<br>prop $= 0.78$ | $\mu_2 = \pi$ (fixed)<br>$\kappa_2 = 6.31$ |

Table A2: AIC and maximum likelihood estimates of parameter values from fitting linear distributions to the step lengths of the fisher data.

| **Distribution** | **AIC** | **Parameters** | |
|---|---|---|---|
| gamma | 61865 | $k = 0.74$ | $\theta = 0.01$ |
| Weibull | 61783 | $k = 0.01$ | $\lambda = 1.61$ |
| lognormal | 61772 | $\mu = 3.83$ | $\sigma = 1.41$ |

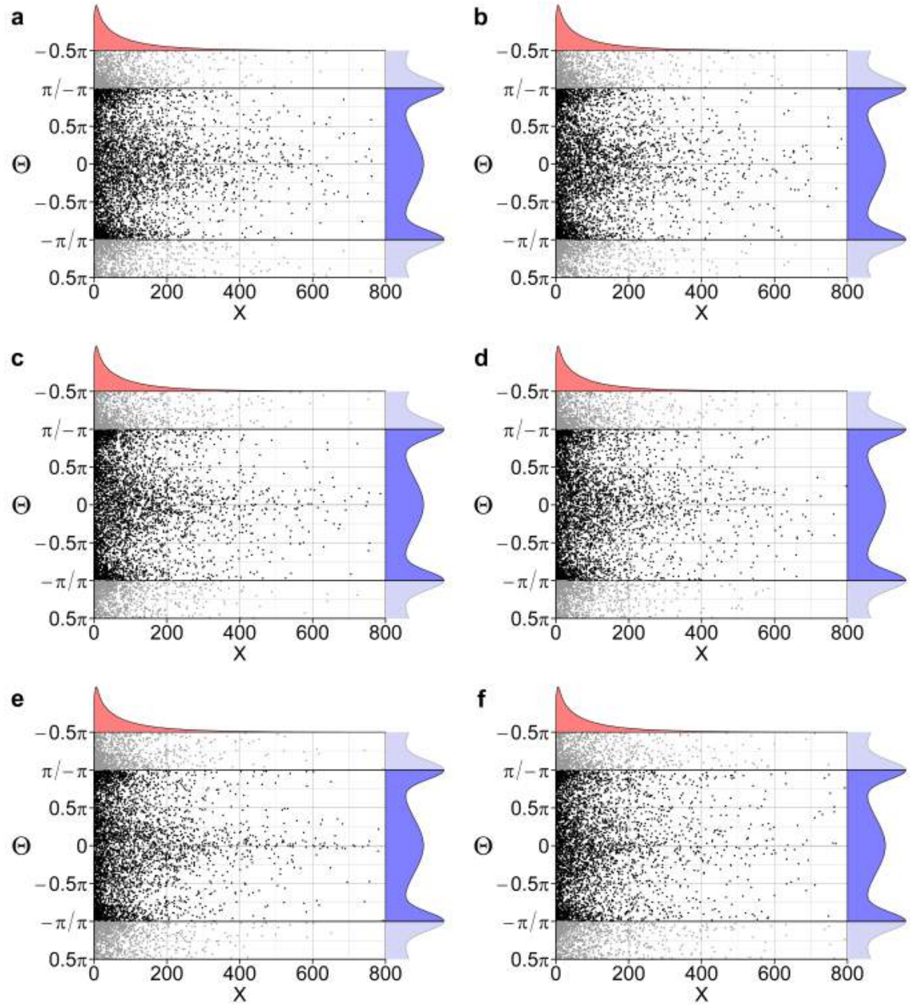


Figure A6: Scatter plots of 4350 step lengths and turn angles sampled from a joint distribution obtained with copulae a-f (see Table 1 in the main text). Maximum likelihood estimates of the marginal densities are plotted in red and blue next to the axes.

##### A4.1 Pseudo-Observations, Copula Samples and Densities

We show below the pseudo-observations of the fisher data and samples from copulae a-f (see main text, table 1). The red line is there as visual aid and is obtained from a least squares regression of a linear model with the linear rank

($s$) as response and sine and cosine of the circular ranks ($r$) as covariates:

$$\mathrm{E}(s_i) = \beta_0 + \beta_1 \cos(r_i) + \beta_2 \sin(r_i) \tag{6}$$

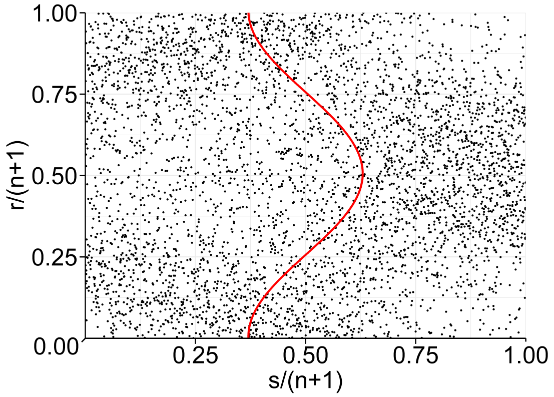


Figure A7: Scatter plot of the 4350 pseudo-observations of the fisher data.

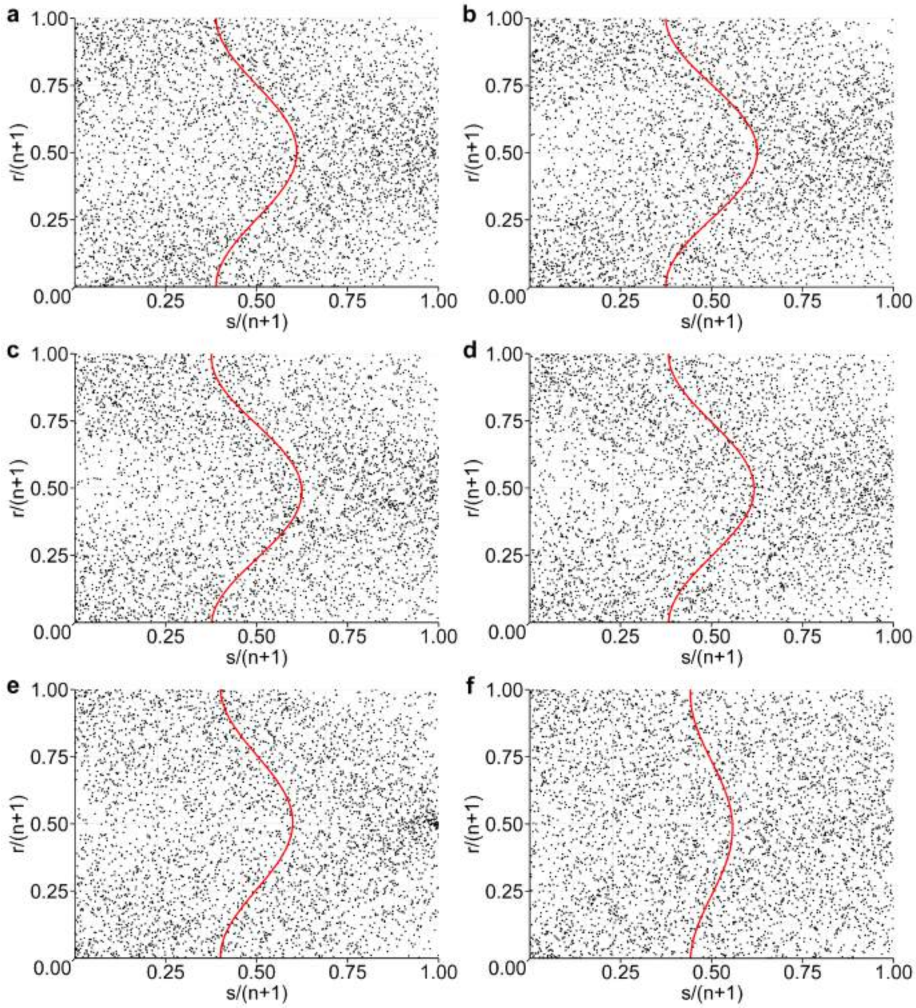


Figure A8: Scatter plot of 4350 samples of copulae a-f (see main text, table 1).

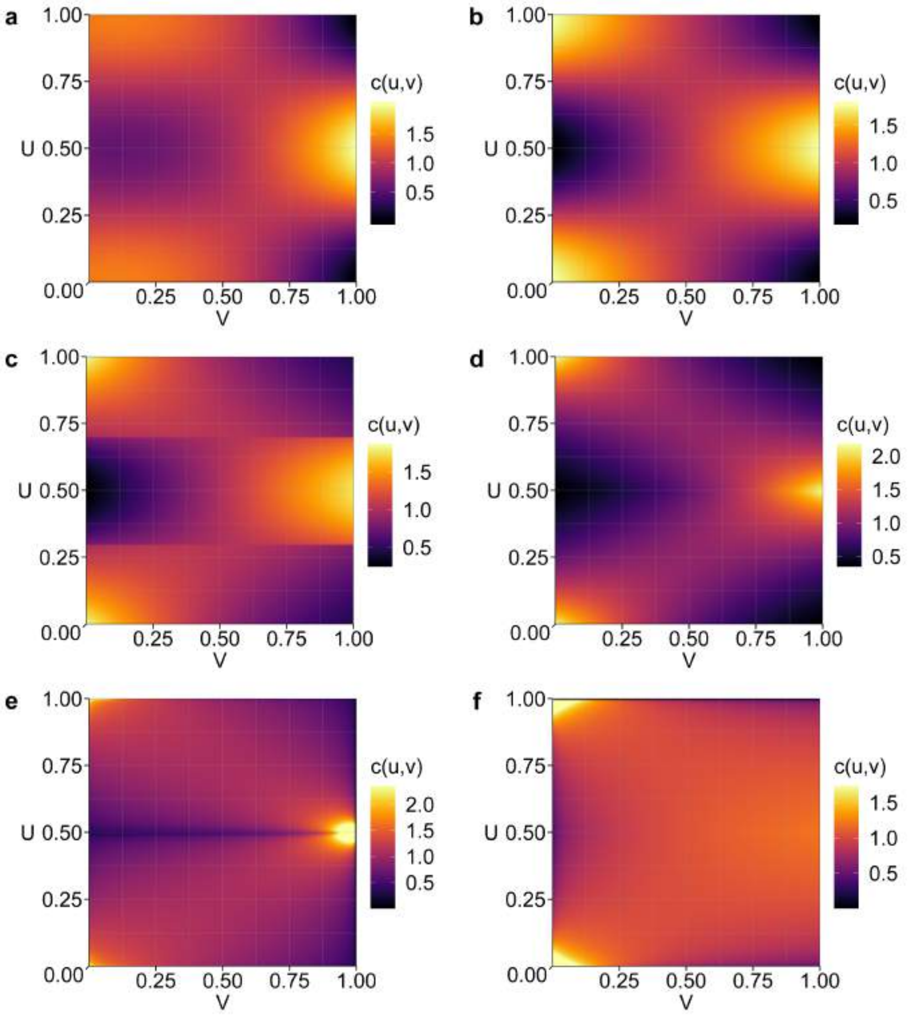


Figure A9: Plots of the copula densities of copulae a-f (see main text, table 1). Note that for copulae e and f, PDF values larger than the 99.5 percentile of all PDF values of that copula are cut off for easier visualization.

#### A4.2 Tracks and Diffusion Coefficients

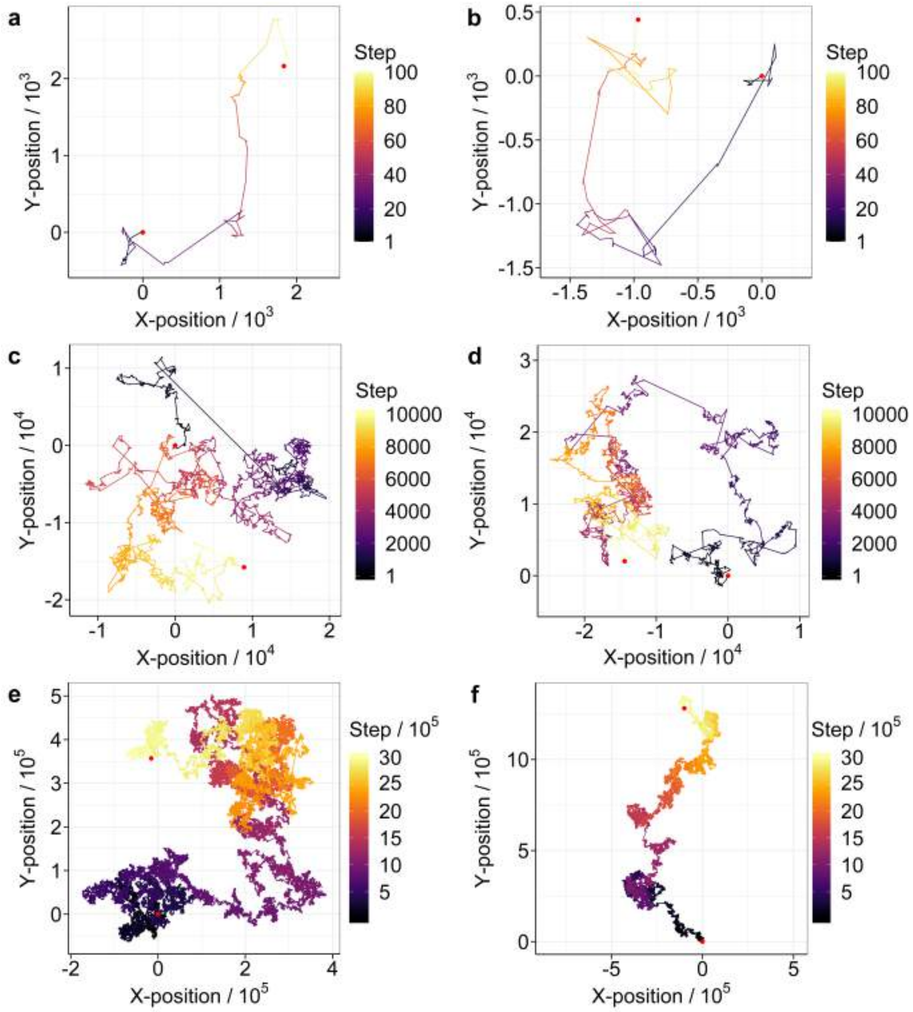


Figure A10: Comparison of tracks of 100, 10,000, and 3,000,000 steps generated from step lengths and turn angles drawn from a joint distribution with correlation structure given by copula a (left column) and with independent marginals (right column). Note that the scales on which the tracks are plotted are not the same for both columns.

The mean square displacement at time $t$ is defined as

$$\text{MSD} \equiv \langle |\mathbf{R}(t) - \mathbf{R}(0)|^2 \rangle \,, \tag{7}$$

where $\mathbf{R}(t)$ is the position of the walker $t$ time steps after some time origin (when the walker was at $\mathbf{R}(0)$). $\langle ... \rangle$ denotes an averaging over walkers and time origins.

From 10 simulated tracks of $3 \times 10^5$ steps, we calculated the MSD for time steps of length 1 to $n_{max} = 1.5 \times 10^5$ and for all possible time origins from 0 to $n_{max}$. In this way, the MSD at every time step is obtained as an average of $10 \times n_{max}$ values. According to Einstein (Einstein 1905), the MSD is proportional to the time step in the limit of infinite time steps. The proportionality constant is the diffusion coefficient, $D$.

$$D = \lim_{t \to \infty} \frac{1}{4t} \langle |\mathbf{R}(t) - \mathbf{R}(0)|^2 \rangle \tag{8}$$

Thus, to determine the diffusion coefficient, we can plot the MSD against the length of the time steps and carry out a least squares regression to find the slope. Excluding smaller time steps in its calculation (when the MSD is not yet in the linear regime), this slope is approximately proportional to the diffusion coefficient. We calculated and plotted the MSD as described above for tracks obtained with copula a and with independent marginals. Visually we determined the limit of the linear regime to be at 100,000 time steps.

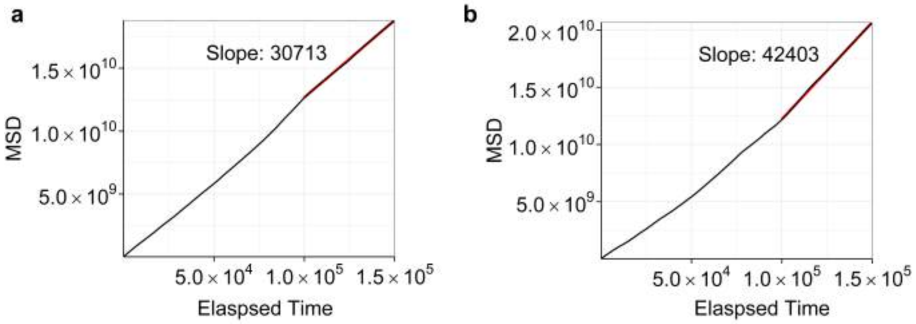


Figure A11: Plots of MSD against time step lengths for tracks obtained with copula a (left) and tracks obtained with independent marginals (right). The least squares regression line is fitted for time steps longer than 80,000 and is shown in red.

With a time step length of 10 minutes (see main text), we can convert the slopes to the diffusion coefficients with SI units, $m^2/s$, by dividing them by $60 \times 10 \times 4 = 2400$ as reported in the main text.
