## Supplementary material for "Circular-Linear Copulae for Animal Movement Data": Code application: Code_application.html

##### July 14, 2021

- Load libraries and data
- Fit marginal distributions
- Fit circular-linear copulae

### Load libraries and data

The code in this section is taken directly from Muff, S., Signer, J., & Fieberg, J. (2019). R code and output supporting: Accounting for individual-specific variation in habitat-selection studies: Efficient estimation of mixed-effects models using Bayesian or frequentist computation. Minneapolis, MN: University of Minnesota Digital Conservancy.

After loading the necessary packages, we extract the important data from the data set. The Fisher movement data was obtained from LaPoint, S., Gallery, P., Wikelski, M. and Kays, R. (2013). Animal behavior, cost-based corridor models, and real corridors. Landscape Ecology, 28, 1615-1630.

```
library(tidyverse)
library(amt)
library(copula)
library(cylcop)
dat <- read.csv("Fisher.csv", header = T) %>%
  filter(!is.na(`location_lat`)) %>%
  select(x = `location_long`,
         y = `location_lat`,
         t = `timestamp`,
         id = `tag_local_identifier`) %>%
  filter(id %in% c(1465, 1466, 1072, 1078, 1016, 1469))
write_csv(dat, "fisher_simple_data.csv")
dat <- read_csv("fisher_simple_data.csv")
dat_all <- dat %>% nest(-id)
dat_all$sex <- c("f", "f", "f", "m", "m", "m")
```

We then create a track from the data and convert it to an accurate projection for that geographic region with unit meters (epsg:5070).

```
dat_all <- dat_all %>%
  mutate(trk = map(data, function(d) {
    make_track(d, x, y, t, crs = sp::CRS("+init=epsg:4326")) %>%
      transform_coords(sp::CRS("+init=epsg:5070"))
  }))
```

Next, we show information on the sampling rate of the different individuals,

```
dat_all %>% mutate(sr = lapply(trk, summarize_sampling_rate)) %>%
  select(id, sr) %>% unnest
```

```
## # A tibble: 6 x 10
##      id   min    q1 median  mean    q3   max    sd     n unit 
##   <dbl> <dbl> <dbl>  <dbl> <dbl> <dbl> <dbl> <dbl> <int> <chr>
## 1  1072 3.92   9.80  10.0  20.7  10.3  1650.  95.2  1348 min  
## 2  1465 0.4    1.97   2.03  8.93  3.98 1080.  48.1  3003 min  
## 3  1466 0.417  1.97   2.07 15.7   4.08 2167. 104.   1500 min  
## 4  1078 1.35   9.78  10.0  21.7  10.3  2111.  87.2  1637 min  
## 5  1469 0.417  1.97   2.17 13.3   5.42 2889.  90.2  2435 min  
## 6  1016 0.100  1.93   2.03  8.04  2.57 1209.  44.0  8957 min
```

resample tracks at a rate based on the sampling rates above,

```
dat_all <- dat_all %>% mutate(dat_clean = map(trk, ~ {
  .x %>% track_resample(rate = minutes(10), tolerance = seconds(60))
}))
```

and convert to steps.

```
dat_all <-
  dat_all %>%
  mutate(stps = map(dat_clean, ~ .x %>% steps_by_burst()))
```

### Fit marginal distributions

For fitting, we will pool together the step lengths and turn angles of all animals. The step length distributions are fitted to all available step lengths, not only to those where the corresponding turning angle is not NA.

```
dat_pool <- dat_all %>% select(stps) %>% unnest()
x <- dat_pool$sl_
theta <- dat_pool$ta_[!is.na(dat_pool$ta_)]
```

We can now fit distributions to the step lengths.

```
cylcop::fit_steplength(x, parametric = "gamma")
cylcop::fit_steplength(x, parametric = "weibull")
marg_dist <- cylcop::fit_steplength(x, parametric = "lognorm")
```

The lowest AIC is achieved with a log-normal distribution.

```
marg_dist
```

```
## $coef
## $coef$meanlog
## [1] 3.832681
## 
## $coef$sdlog
## [1] 1.412453
## 
## 
## $se
## $se$meanlog
## [1] 0.01901443
## 
## $se$sdlog
## [1] 0.01344523
## 
## 
## $logL
## [1] -30883.95
## 
## $AIC
## [1] 61771.91
## 
## $name
## [1] "lnorm"
```

Next, we fit circular distributions to the turn angles.

```
cylcop::fit_angle(theta, parametric = "mixedvonmises")
cylcop::fit_angle(theta, parametric = "vonmises")
cylcop::fit_angle(theta, parametric = "wrappedcauchy")
marg_angle <- cylcop::fit_angle(theta, parametric = "mixedvonmises", mu = c(0, pi))
```

The lowest AIC is achieved with a mixed von Mises distribution with fixed location parameters.

```
marg_angle
```

```
## $coef
## $coef$mu1
## [1] 0
## 
## $coef$mu2
## [1] 3.141593
## 
## $coef$kappa1
## [1] 0.4839511
## 
## $coef$kappa2
## [1] 6.307059
## 
## $coef$prop
## [1] 0.7757561
## 
## 
## $se
## list()
## 
## $logL
## [1] -7791.917
## 
## $AIC
## [1] 15589.83
## 
## $name
## [1] "mixedvonmises"
```

### Fit circular-linear copulae

First, we remove step lengths that correspond to NA in turn angles from the data.

```
x <- dat_pool$sl_[!is.na(dat_pool$ta_)]
```

Next, we investigate circular-linear correlation measures of the bivariate data.

```
cylcop::cor_cyl(theta = theta, x = x)
```

```
## [1] 0.1024313
```

```
cylcop::mi_cyl(
  theta = theta,
  x = x,
  normalize = T,
  symmetrize = T
)
```

```
## [1] 0.03732642
```

After this, we roughly estimate copula parameters using a grid search.

```
start_quadsec <- 
  cylcop::optCor(
  copula = cyl_quadsec(),
  theta = theta,
  x = x,
  method = "cor_cyl"
)
start_quadsec <-
  cylcop::optCor(
    copula = cyl_quadsec(),
    theta = theta,
    x = x,
    method = "mi_cyl"
  )
start_cubsec <-
  cylcop::optCor(
    copula = cyl_cubsec(),
    theta = theta,
    x = x,
    method = "cor_cyl",
    acc = 0.05,
    n = 10000
  )
```

For cyl\_rect\_combine copulae with symmetric rectangles spanning the entire unit square, we can calculate parameters from Kendall’s tau.

```
start_rect_frank <-
  cylcop::optCor(
    copula = cyl_rect_combine(
      copula = frankCopula(),
      low_rect = c(0, 0.5),
      up_rect = "symmetric"
    ),
    theta = theta,
    x = x,
    method = "tau"
  )
start_rect_clayton <-
  cylcop::optCor(
    copula = cyl_rect_combine(
      copula = claytonCopula(),
      low_rect = c(0, 0.5),
      up_rect = "symmetric"
    ),
    theta = theta,
    x = x,
    method = "tau"
  )
start_rect_gumbel <-
  cylcop::optCor(
    copula = cyl_rect_combine(
      copula = gumbelCopula(),
      low_rect = c(0, 0.5),
      up_rect = "symmetric"
    ),
    theta = theta,
    x = x,
    method = "tau"
  )
```

Using these parameter values as starting points, we optimize copula parameters with a maximum pseudo likelihood approach.

```
copula_quadsec <-
  cylcop::optML(
    copula = cyl_quadsec(),
    theta = theta,
    x = x,
    parameters = "a",
    start = start_quadsec
  )

copula_cubsec <-
  cylcop::optML(
    copula = cyl_cubsec(),
    theta = theta,
    x = x,
    parameters = c("a", "b"),
    start = start_cubsec,
    traceOpt = T
  )

copula_rect_frank <-
  cylcop::optML(
    copula = cyl_rect_combine(
      copula = frankCopula(param = 2),
      low_rect = c(0, 0.5),
      up_rect = "symmetric"
    ),
    theta = theta,
    x = x,
    parameters = "alpha",
    start = start_rect_frank,
    traceOpt = T
  )
```

With a Clayton copula with negative parameter the rectangular patchwork copula fits the data very badly, and we therefore set the lower bound close to 0.

```
 copula_rect_clayton <-
   cylcop::optML(
     copula = cyl_rect_combine(
       copula = claytonCopula(),
       low_rect = c(0, 0.5),
       up_rect = "symmetric"
     ),
     theta = theta,
     x = x,
     parameters = "alpha",
     start = start_rect_clayton,
     traceOpt = T,
     lower = 0.0001
   )
 copula_rect_gumbel <-
   cylcop::optML(
     copula = cyl_rect_combine(
       copula = gumbelCopula(),
       low_rect = c(0, 0.5),
       up_rect = "symmetric"
     ),
     theta = theta,
     x = x,
     parameters = "alpha",
     start = start_rect_gumbel,
     traceOpt = T
   )
```

With the following copula below, optimizing the upper bound of the lower rectangle (and with it the lower bound of the upper rectangle, because up\_rect=“symmetric”) results either in rectangles of area 0 and a cyl\_quadsec copula or in rectangles spanning the entire unit square, i.e. copula\_rect\_frank.

```
 cylcop::optML(
   copula = cyl_rect_combine(
     copula = frankCopula(param = 2),
     low_rect = c(0, 0.5),
     up_rect = "symmetric",
     background = cyl_quadsec()
   ),
   theta = theta,
   x = x,
   parameters = c("alpha", "low_rect2", "bg_a"),
   start = c(1.415, 0.357, 0.113),
   lower = c(0.2, 0, 0),
   upper = c(10, 0.45, 1 / (2 * pi)),
   traceOpt = T
 )
```

Instead, we arbitrarily set the rectangles to R\_1=[0.0,0.3]x[0,1] and R\_2=[0.7,1.0]x[0,1]

```
 copula_rect_frank_quadsec <- cylcop::optML(
   copula = cyl_rect_combine(
     copula = frankCopula(param = 2),
     low_rect = c(0, 0.3),
     up_rect = "symmetric",
     background = cyl_quadsec()
   ),
   theta = theta,
   x = x,
   parameters = c("alpha", "bg_a"),
   start = c(1.415, 0.113),
   lower = c(0.0001, 0),
   traceOpt = T
 )
```

The copula giving the lowest AIC score was the cubic sections copula.

```
copula_cubsec
```

```
## $copula
## Cub. sect. copula 
## a = 0.06604284 
## b = -0.1591549 
## 
## $logL
## [1] 274.8908
## 
## $AIC
## [1] -545.7815
```

```
sessionInfo()
```

```
## R version 4.0.5 (2021-03-31)
## Platform: x86_64-w64-mingw32/x64 (64-bit)
## Running under: Windows 10 x64 (build 19042)
## 
## Matrix products: default
## 
## locale:
## [1] LC_COLLATE=German_Switzerland.1252  LC_CTYPE=German_Switzerland.1252   
## [3] LC_MONETARY=German_Switzerland.1252 LC_NUMERIC=C                       
## [5] LC_TIME=German_Switzerland.1252    
## 
## attached base packages:
## [1] stats     graphics  grDevices utils     datasets  methods   base     
## 
## other attached packages:
##  [1] cylcop_0.0.0.9000 copula_1.0-1      amt_0.1.4         forcats_0.5.1    
##  [5] stringr_1.4.0     dplyr_1.0.5       purrr_0.3.4       readr_1.4.0      
##  [9] tidyr_1.1.3       tibble_3.1.1      ggplot2_3.3.3     tidyverse_1.3.1  
## 
## loaded via a namespace (and not attached):
##   [1] colorspace_2.0-0        ellipsis_0.3.1          rgdal_1.5-23           
##   [4] fs_1.5.0                rstudioapi_0.13         circular_0.4-93        
##   [7] gsl_2.1-6               fansi_0.4.2             mvtnorm_1.1-1          
##  [10] lubridate_1.7.10        xml2_1.3.2              codetools_0.2-18       
##  [13] splines_4.0.5           knitr_1.33              jsonlite_1.7.2         
##  [16] broom_0.7.6             dbplyr_2.1.1            stabledist_0.7-1       
##  [19] GoFKernel_2.1-1         shiny_1.6.0             compiler_4.0.5         
##  [22] httr_1.4.2              backports_1.2.1         assertthat_0.2.1       
##  [25] Matrix_1.3-2            fastmap_1.1.0           lazyeval_0.2.2         
##  [28] cli_2.5.0               later_1.2.0             htmltools_0.5.1.1      
##  [31] tools_4.0.5             gtable_0.3.0            glue_1.4.2             
##  [34] movMF_0.2-5             Rcpp_1.0.6              slam_0.1-48            
##  [37] cellranger_1.1.0        jquerylib_0.1.4         raster_3.4-5           
##  [40] vctrs_0.3.7             crosstalk_1.1.1         xfun_0.22              
##  [43] rbibutils_2.1.1         rvest_1.0.0             mime_0.10              
##  [46] miniUI_0.1.1.1          lifecycle_1.0.0         MASS_7.3-53.1          
##  [49] scales_1.1.1            hms_1.0.0               promises_1.2.0.1       
##  [52] yaml_2.2.1              gridExtra_2.3           sass_0.3.1             
##  [55] stringi_1.5.3           highr_0.9               pcaPP_1.9-74           
##  [58] boot_1.3-27             manipulateWidget_0.10.1 Rdpack_2.1.1           
##  [61] rlang_0.4.10            pkgconfig_2.0.3         rgl_0.106.8            
##  [64] evaluate_0.14           lattice_0.20-41         htmlwidgets_1.5.3      
##  [67] tidyselect_1.1.0        magrittr_2.0.1          R6_2.5.0               
##  [70] generics_0.1.0          ADGofTest_0.3           DBI_1.1.1              
##  [73] pillar_1.6.0            haven_2.4.1             withr_2.4.2            
##  [76] survival_3.2-11         sp_1.4-5                pspline_1.0-18         
##  [79] modelr_0.1.8            crayon_1.4.1            KernSmooth_2.23-18     
##  [82] utf8_1.2.1              plotly_4.9.3            rmarkdown_2.7          
##  [85] viridis_0.6.0           grid_4.0.5              readxl_1.3.1           
##  [88] data.table_1.14.0       infotheo_1.2.0          reprex_2.0.0           
##  [91] digest_0.6.27           webshot_0.5.2           xtable_1.8-4           
##  [94] httpuv_1.6.0            extraDistr_1.9.1        numDeriv_2016.8-1.1    
##  [97] stats4_4.0.5            munsell_0.5.0           viridisLite_0.4.0      
## [100] bslib_0.2.4
```
