## Supplementary material for "Circular-Linear Copulae for Animal Movement Data": Code application: Code_application.R

```
#'---
#'title: "Circular-Linear Copulae for Animal Movement Data"  
#'author: "Florian Hodel and John Fieberg"
#'date: "July 14, 2021"
#'output:
#'  html_document:
#'    toc: yes
#'---
#'
## ----setup, include=FALSE-------------------------------------------------------------
knitr::opts_chunk$set(echo = TRUE)

#' ## Load libraries and data
#' The code in this section is taken directly from Muff,
#' S., Signer, J., & Fieberg, J. (2019). R code and output supporting:
#' Accounting for individual-specific variation in habitat-selection studies:
#' Efficient estimation of mixed-effects models using Bayesian or
#' frequentist computation. Minneapolis, MN: University of Minnesota Digital
#' Conservancy.
#' 
#' After loading the necessary packages, we extract the important
#' data from the data set. The Fisher movement data was obtained from
#' LaPoint, S., Gallery, P., Wikelski, M. and Kays, R. (2013).
#' Animal behavior, cost-based corridor models, and real corridors.
#' Landscape Ecology, 28, 1615-1630.
## ---- eval=T, echo=T,warning=FALSE,message=FALSE,results='hide'-----------------------
library(tidyverse)
library(amt)
library(copula)
library(cylcop)
dat <- read.csv("Fisher.csv", header = T) %>%
  filter(!is.na(`location_lat`)) %>%
  select(x = `location_long`,
         y = `location_lat`,
         t = `timestamp`,
         id = `tag_local_identifier`) %>%
  filter(id %in% c(1465, 1466, 1072, 1078, 1016, 1469))
write_csv(dat, "fisher_simple_data.csv")
dat <- read_csv("fisher_simple_data.csv")
dat_all <- dat %>% nest(-id)
dat_all$sex <- c("f", "f", "f", "m", "m", "m")

#'We then create a track from the data and convert it to an accurate projection
#' for that geographic region with unit meters (epsg:5070).
## ---- eval=T, echo=T,warning=FALSE,message=FALSE,results='hide'-----------------------
dat_all <- dat_all %>%
  mutate(trk = map(data, function(d) {
    make_track(d, x, y, t, crs = sp::CRS("+init=epsg:4326")) %>%
      transform_coords(sp::CRS("+init=epsg:5070"))
  }))

#' Next, we show information on the sampling rate of the different individuals,
## ---- eval=T, echo=T,warning=FALSE,message=FALSE--------------------------------------
dat_all %>% mutate(sr = lapply(trk, summarize_sampling_rate)) %>%
  select(id, sr) %>% unnest

#' resample tracks at a rate based on the sampling rates above,
## ---- eval=T, echo=T,warning=FALSE,message=FALSE,results='hide'-----------------------
dat_all <- dat_all %>% mutate(dat_clean = map(trk, ~ {
  .x %>% track_resample(rate = minutes(10), tolerance = seconds(60))
}))

#' and convert to steps.
## ---- eval=T, echo=T,warning=FALSE,message=FALSE,results='hide'-----------------------
dat_all <-
  dat_all %>%
  mutate(stps = map(dat_clean, ~ .x %>% steps_by_burst()))


#' ## Fit marginal distributions
#' 
#' For fitting, we will pool together the step lengths and turn angles of all animals.
#' The step length distributions are fitted to all available step lengths,
#' not only to those where the corresponding turning angle is not NA.
## ---- eval=T, echo=T,warning=FALSE,message=FALSE,results='hide'-----------------------
dat_pool <- dat_all %>% select(stps) %>% unnest()
x <- dat_pool$sl_
theta <- dat_pool$ta_[!is.na(dat_pool$ta_)]

#' We can now fit distributions to the step lengths.
## ---- eval=T, echo=T,warning=FALSE,message=FALSE,results='hide'-----------------------
cylcop::fit_steplength(x, parametric = "gamma")
cylcop::fit_steplength(x, parametric = "weibull")
marg_dist <- cylcop::fit_steplength(x, parametric = "lognorm")

#' The lowest AIC is achieved with a log-normal distribution.
## ---- eval=T, echo=T,warning=FALSE,message=FALSE--------------------------------------
marg_dist

#' Next, we fit circular distributions to the turn angles.
## ---- eval=T, echo=T,warning=FALSE,message=FALSE,results='hide'-----------------------
cylcop::fit_angle(theta, parametric = "mixedvonmises")
cylcop::fit_angle(theta, parametric = "vonmises")
cylcop::fit_angle(theta, parametric = "wrappedcauchy")
marg_angle <- cylcop::fit_angle(theta, parametric = "mixedvonmises", mu = c(0, pi))

#' The lowest AIC is achieved with a mixed von Mises distribution with fixed
#' location parameters.
## ---- eval=T, echo=T,warning=FALSE,message=FALSE--------------------------------------
marg_angle

#' ## Fit circular-linear copulae
#' First, we remove step lengths that correspond to NA in turn angles
#' from the data.
## ---- eval=T, echo=T,warning=FALSE,message=FALSE,results='hide'-----------------------
x <- dat_pool$sl_[!is.na(dat_pool$ta_)]

#' Next, we investigate circular-linear correlation measures of
#' the bivariate data.
## ---- eval=T, echo=T,warning=FALSE,message=FALSE--------------------------------------
cylcop::cor_cyl(theta = theta, x = x)
cylcop::mi_cyl(
  theta = theta,
  x = x,
  normalize = T,
  symmetrize = T
)

#' After this, we roughly estimate copula parameters using a grid search.
## ---- eval=T, echo=T,warning=FALSE,message=FALSE,results='hide'-----------------------
start_quadsec <- 
  cylcop::optCor(
  copula = cyl_quadsec(),
  theta = theta,
  x = x,
  method = "cor_cyl"
)
start_quadsec <-
  cylcop::optCor(
    copula = cyl_quadsec(),
    theta = theta,
    x = x,
    method = "mi_cyl"
  )
start_cubsec <-
  cylcop::optCor(
    copula = cyl_cubsec(),
    theta = theta,
    x = x,
    method = "cor_cyl",
    acc = 0.05,
    n = 10000
  )

#' For cyl_rect_combine copulae with symmetric rectangles spanning the entire
#' unit square, we can calculate parameters from Kendall's tau.
## ---- eval=T, echo=T,warning=FALSE,message=FALSE,results='hide'-----------------------
start_rect_frank <-
  cylcop::optCor(
    copula = cyl_rect_combine(
      copula = frankCopula(),
      low_rect = c(0, 0.5),
      up_rect = "symmetric"
    ),
    theta = theta,
    x = x,
    method = "tau"
  )
start_rect_clayton <-
  cylcop::optCor(
    copula = cyl_rect_combine(
      copula = claytonCopula(),
      low_rect = c(0, 0.5),
      up_rect = "symmetric"
    ),
    theta = theta,
    x = x,
    method = "tau"
  )
start_rect_gumbel <-
  cylcop::optCor(
    copula = cyl_rect_combine(
      copula = gumbelCopula(),
      low_rect = c(0, 0.5),
      up_rect = "symmetric"
    ),
    theta = theta,
    x = x,
    method = "tau"
  )

#' Using these parameter values as starting points, we optimize copula
#' parameters with a maximum pseudo likelihood approach.
## ---- eval=T, echo=T,warning=FALSE,message=FALSE,results='hide'-----------------------
copula_quadsec <-
  cylcop::optML(
    copula = cyl_quadsec(),
    theta = theta,
    x = x,
    parameters = "a",
    start = start_quadsec
  )

copula_cubsec <-
  cylcop::optML(
    copula = cyl_cubsec(),
    theta = theta,
    x = x,
    parameters = c("a", "b"),
    start = start_cubsec,
    traceOpt = T
  )

copula_rect_frank <-
  cylcop::optML(
    copula = cyl_rect_combine(
      copula = frankCopula(param = 2),
      low_rect = c(0, 0.5),
      up_rect = "symmetric"
    ),
    theta = theta,
    x = x,
    parameters = "alpha",
    start = start_rect_frank,
    traceOpt = T
  )

#' With a Clayton copula with negative parameter the rectangular patchwork
#' copula fits the data very badly, and we therefore set the lower
#' bound close to 0.
## ---- eval=F, echo=T,warning=FALSE,message=FALSE,results='hide'-----------------------
 copula_rect_clayton <-
   cylcop::optML(
     copula = cyl_rect_combine(
       copula = claytonCopula(),
       low_rect = c(0, 0.5),
       up_rect = "symmetric"
     ),
     theta = theta,
     x = x,
     parameters = "alpha",
     start = start_rect_clayton,
     traceOpt = T,
     lower = 0.0001
   )
 copula_rect_gumbel <-
   cylcop::optML(
     copula = cyl_rect_combine(
       copula = gumbelCopula(),
       low_rect = c(0, 0.5),
       up_rect = "symmetric"
     ),
     theta = theta,
     x = x,
     parameters = "alpha",
     start = start_rect_gumbel,
     traceOpt = T
   )

#' With the following copula below, optimizing the upper bound of the lower
#' rectangle (and with it the lower bound of the upper rectangle, because
#' up_rect="symmetric") results either in rectangles of area 0 and a cyl_quadsec
#' copula or in rectangles spanning the entire unit square,
#' i.e. copula_rect_frank.
## ---- eval=F, echo=T,warning=FALSE,message=FALSE,results='hide'-----------------------
 cylcop::optML(
   copula = cyl_rect_combine(
     copula = frankCopula(param = 2),
     low_rect = c(0, 0.5),
     up_rect = "symmetric",
     background = cyl_quadsec()
   ),
   theta = theta,
   x = x,
   parameters = c("alpha", "low_rect2", "bg_a"),
   start = c(1.415, 0.357, 0.113),
   lower = c(0.2, 0, 0),
   upper = c(10, 0.45, 1 / (2 * pi)),
   traceOpt = T
 )

#' Instead, we arbitrarily set the rectangles to R_1=[0.0,0.3]x[0,1]
#' and R_2=[0.7,1.0]x[0,1]
## ---- eval=F, echo=T,warning=FALSE,message=FALSE,results='hide'-----------------------
 copula_rect_frank_quadsec <- cylcop::optML(
   copula = cyl_rect_combine(
     copula = frankCopula(param = 2),
     low_rect = c(0, 0.3),
     up_rect = "symmetric",
     background = cyl_quadsec()
   ),
   theta = theta,
   x = x,
   parameters = c("alpha", "bg_a"),
   start = c(1.415, 0.113),
   lower = c(0.0001, 0),
   traceOpt = T
 )

#' The copula giving the lowest AIC score was the cubic sections copula.
## ---- eval=T, echo=T,warning=FALSE,message=FALSE--------------------------------------
copula_cubsec

## ---- eval=T, echo=T,warning=FALSE,message=FALSE--------------------------------------
sessionInfo()
```
